# The endophytic fungus *Serendipita indica* alters auxin distribution in *Arabidopsis thaliana* roots through alteration of auxin transport and conjugation to promote plant growth

**DOI:** 10.1101/2024.01.23.576781

**Authors:** Adrián González Ortega-Villaizán, Eoghan King, Manish K. Patel, Marta-Marina Pérez- Alonso, Sandra S. Scholz, Hitoshi Sakakibara, Takatoshi Kiba, Mikiko Kojima, Yumiko Takebayashi, Patricio Ramos, Luis Morales-Quintana, Sarah Breitenbach, Ana Smolko, Branka Salopek-Sondi, Nataša Bauer, Jutta Ludwig-Müller, Anne Krapp, Ralf Oelmüller, Jesús Vicente-Carbajosa, Stephan Pollmann

## Abstract

Plants share their habitats with a multitude of different microbes. This close vicinity promoted the evolution of inter-organismic interactions between plants and many different microorganisms that provide mutual growth benefits both to the plant and the microbial partner. The symbiosis of *Arabidopsis thaliana* with the beneficial root colonizing endophyte *Serendipita indica* represents a well-studied system. Co-colonization of Arabidopsis roots with *S. indica* significantly promotes plant growth. Due to the notable phenotypic alterations of fungus-infected root systems, the involvement of a reprogramming of plant hormone levels, especially that of indole-3-acetic acid, has been suggested earlier. However, until now, the molecular mechanism by which *S. indica* promotes plant growth remains largely unknown. This study used comprehensive transcriptomics, metabolomics, reverse genetics, and life cell imaging to reveal the intricacies of auxin-related processes that affect root growth in the symbiosis between *A. thaliana* and *S. indica*. Our experiments revealed the essential role of tightly controlled auxin conjugation in the plant–fungus interaction. It particularly highlighted the importance of two *GRETCHEN HAGEN 3* (*GH3*) genes, *GH3.5* and *GH3.17*, for the fungus infection-triggered stimulation of biomass production, thus broadening our knowledge about the function of GH3s in plants. Furthermore, we provide evidence for the transcriptional alteration of the *PIN2* auxin transporter gene in roots of Arabidopsis seedlings infected with *S. indica* and demonstrate that this transcriptional adjustment affects auxin signaling in roots, which results in increased plant growth.

## 1. Introduction

The current global climate change scenario poses a remarkable thread to contemporary agriculture, which is further aggravated by the previsions on the development of the world population over the next decades, which together puts food security into jeopardy (van Dijk et al., 2021). Given that technical progress in agriculture is not expected to provide more than a 20 to 25% increase in crop production in the near future, alternative and possibly more holistic approaches need to be embraced to address the problem of food security.

The application of beneficial microbial symbionts as biofertilizers to boost crop productivity might be a solution to considerably improve agricultural productivity in a sustainable manner (Pérez-Alonso et al., 2020). There are numerous examples of growth-promoting microbes in the literature and how abiotic stresses affect the interaction between plants and microbes has just recently been reviewed (Bastías et al., 2022). However, to fully exploit the described interactions between plants and their symbionts, it is paramount to understand the underlying molecular mechanisms that build the framework of these interactions.

A good example of a beneficial interaction between plants and their symbionts is the interaction between the root colonizing endophyte *Serendipita indica* (formerly called *Piriformospora indica*) and its wide range of host plants, including several important crops such as barley, wheat, and corn (Hosseini et al., 2017; Mensah et al., 2020; Waller et al., 2005; Zhang et al., 2018). *S. indica* is an axenically cultivable root endophyte of the order Sebacinales (Weiss et al., 2016) that promotes plant performance, biomass production, and tolerance to a wide array of biotic and abiotic stresses (Jogawat et al., 2016; Peškan-Berghöfer et al., 2004; Sun et al., 2014; Varma et al., 1999). However, despite a large body of evidence that describes different facets of the physiological impact of the interaction of *S. indica* with its host plants, our current understanding of the molecular mechanisms involved in the establishment and maintenance of the symbiotic interaction is still elusive. Apart from a large body of previous studies addressing processes associated with the initiation of the plant–fungus interaction, which later merges into the limitation of endophyte proliferation in root tissue (Jacobs et al., 2011; Lahrmann et al., 2015; Zuccaro et al., 2011), little is known about the molecular implications that trigger lingering plant growth in response to an infection with *S. indica*. However, strict control of the contents of different plant hormones and other small signaling molecules, such as cytosolic calcium, have been reported to play crucial roles (Nongbri et al., 2012; Pérez-Alonso et al., 2022; Vadassery et al., 2008; Xu et al., 2018).

With respect to the control of root growth, the tight regulation of auxin fluxes and the formation of local maxima across the developing root are pivotal (Petrášek & Friml, 2009; Roychoudhry & Kepinski, 2022). In addition to the importance of polar auxin transport for proper root development, there is mounting evidence that local auxin biosynthesis is also crucial to control root growth and development in plants (Brumos et al., 2018). However, much less is known about the role of auxin degradation and indole-3-acetic acid (IAA) sequestration through the formation of sugar or amino acid conjugates, respectively (Casanova-Sáez et al., 2022; Mateo-Bonmatí et al., 2021; Mellor et al., 2016; Porco et al., 2016). In the context of plant–microbe interactions, the conjugation of free IAA with amino acids catalyzed by auxin-inducible acyl amino synthetases of group II of GRETCHEN HAGEN 3 (GH3) enzymes appears to play an important role, as several studies already demonstrated induction of *GH3* gene expression after microbe infection (Jahn et al., 2013; Wojtaczka et al., 2022; Zhang et al., 2007; Zhang et al., 2008).

In this study, we identified the acyl amido synthetases GH3.5 and GH3.17 as key molecular components that contribute to the establishment and maintenance of the mutual interaction between *Arabidopsis thaliana* and *S. indica*. Transcriptomics analysis revealed the differential expression of several *GH3* genes in fungus-infected Arabidopsis plants. Consistent with an induction of auxin conjugating enzymes upon *S. indica*-infection, we were able to demonstrate that the co-cultivation of an auxin overproducing mutant with the fungus was sufficient to restore a wild type-like phenotype of the mutant. Subsequent RNA-seq experiments underlined the increased expression of *GH3* genes under these conditions. Further reverse genetics studies, employing several *gh3* mutant lines and auxin conjugate hydrolase overexpressing lines (*35S::IAR3*), respectively, accompanied by confocal laser scanning microscopy corroborated our hypothesis that locally restricted auxin conjugation is an essential asset for the mutual interaction between Arabidopsis and *S. indica*. Furthermore, we provide evidence for the repression of the auxin exporter *PIN2* in *S. indica*-infected Arabidopsis roots, which is suggested to translate into local auxin maxima in the root tips. Taken together, our results establish the intimate interplay between local auxin accumulation through *PIN2* repression and the subsequent conjugation of free IAA by the activity of GH3.5 and GH3.17 as key requirements for the successful creation of the symbiosis between *A. thaliana* and the root colonizing endophytic fungus *S. indica*. At the same time, this alteration of the interplay between auxin transport and conjugation in root tips was shown to contribute to the observed fungus-triggered promotion of biomass production.

## 2. Materials and Methods

### 2.1 Biological material and growth conditions

In the presented study, wild-type *Arabidopsis thaliana* (Col-0, stock N1092 and Ler-0, stock NW20), and several previously described mutant lines were used, including the single T-DNA insertion mutants *gh3.4* (Jahn et al., 2013), *gh3.5* and *gh3.17* (Staswick et al., 2005), as well as the EMS mutants *agr1-1* (stock N268) and *agr1-2* (stock N269) (Bell & Maher, 1990; Chen et al., 1998). In addition, the auxin overproduction line YUC9ox (Hentrich et al., 2013), the reporter lines *DR5::Luciferase* (*DR5::Luc*) (Moreno-Risueno et al., 2010), *pGH3.5::3*×*YFP* and *pGH3.17::3*×*YFP* (Pierdonati et al., 2019), as well as *pPIN2::PIN2-GFP* (Xu & Scheres, 2005) were used. After stratification (2 days, 4 °C), plants were grown sterilely on solidified 0.5× MS medium supplemented with 1% (w/v) sucrose (Murashige & Skoog, 1962). Plants were grown in growth chambers under strictly controlled environmental conditions (16 h light, 8 h darkness, constant temperature of 22 °C, 100 to 105 µmol photons m^-2^ s^-1^ photosynthetically active radiation). Furthermore, the beneficial root endophyte fungus *Serendipita indica* strain DSM 11827, which was obtained from the German Collection of Microorganisms and Cell Cultures (DSZM) in Braunschweig, Germany, was used. The fungus was grown in darkness at a constant temperature of 28 °C on solidified arginine phosphate (AP) medium (Rodríguez-Navarro & Ramos, 1984). The fungus was weekly refreshed.

### 2.2 Generation of transgenic Arabidopsis plants

The reporter lines *pGH3.5::Luc* and *pGH3.17::Luc* were generated by amplifying the 2 kb fragments upstream of the start codon of the *GH3.5* (At4g27260) and *GH3.17* (At1g28130) genes by PCR from genomic DNA extracted from *A. thaliana* seedlings. The obtained DNA fragments for the promoters of *GH3.5* and *GH3.17*, respectively, were integrated into the pENTR^TM^/D-TOPO^TM^ vector using the pENTR^TM^/D-TOPO^TM^ cloning kit (Thermo Fisher), and subsequently transferred into the pGWB435 Gateway-compatible destination vector (Nakagawa et al., 2007) to generate the *pGH3.5::Luc* and *pGH3.17::Luc* vector constructs.

For the cloning of the Arabidopsis *IAR3* (At1g51760) coding sequence, total RNA was extracted from one week-old Arabidopsis seedlings and reverse transcribed into cDNA using M-MLV reverse transcriptase (Promega) according to the manufacturer’s instructions. Subsequently, the cDNA was PCR-amplified using the primer pair AtIAR3_*att*B1 and AtIAR3_*att*B2, thereby incorporating a His_6_-tag before the stop codon. The resulting fragment was introduced into the vector pDONR221 (Thermo Scientific) by carrying out a BP reaction, before the modified cDNA fragment was introduced into the *att*R sites of the binary vector pMDC32 (Curtis & Grossniklaus, 2003) by an LR clonase reaction.

Sequence integrity of the different constructs was confirmed by commercial sequencing. The resulting vectors were transformed into *A. thaliana* (Col-0) plants using the Agrobacterium-mediated floral dip method (Clough & Bent, 1998). Independent transgenic lines were selected on solidified 0.5× MS medium plates containing 50 µg/ml kanamycin (pGWB435 derivatives) and 50 µg/ml hygromycin (pMDC32 derivative), respectively. Plants were selfed and selected to homozygosity. Transgenic lines were tested by PCR for the presence of the transgene. All corresponding primer sequences can be found in **Supporting Information:Data Sheet 1**. Additionally, the correct expression of the transgenes was examined. *GH3* genes are known to show a strong auxin response (Hagen & Guilfoyle, 2002). Thus, to select for suitable *GH3*-*Luc* constructs, plants were sprayed with 100 µM indole-3-acetic acid (IAA) and then tested for bioluminescence. The expression of the 35S-driven *IAR3* gene was monitored by protein immunodetection with anti-His (Roche) and anti-mouse IgG-peroxidase (Sigma Aldrich) antibodies. Homozygous T_3_ plants were used for all further experiments.

### 2.3 Root growth assay

To investigate differences in *S. indica*-triggered root growth promotion in the different Arabidopsis genotypes, surface-sterilized seeds were plated on square 0.5× MS plates. Following a stratification of two days at 4 °C, the plates were transferred to a growth chamber and the seedlings were grown vertically for seven days. Thereafter, four seedlings were transferred to fresh square Petri dishes containing solidified Plant Nutrition Medium (PNM) (Johnson et al., 2013). Each seedling was then associated with a 5 mm Ø medium plug extracted from either sterile AP plates (control) or from AP plates harboring a one-week-old *S. indica* mycelium (co-cultivation). Alternatively, the seedlings of each plate were inoculated with 50 µl of a solution containing 2 × 10^5^ spores ml^-1^. The resulting PNM plates were transferred to a growth chamber and maintained at 22-24 °C, 16/8 h photoperiod, 100 µmol photons m^-2^ s^-1^ light intensity for another 10 days. Subsequently, the plants were photographed for the further analysis of the root system.

### 2.4 Quantification of root system architecture traits

The root system of control and *S. indica*-infected Arabidopsis seedlings was captured with a digital camera at a fixed distance (29 cm). Using Adobe Photoshop, the images were cropped to a height of 14 cm keeping only the part comprising the root system. Next, the images were converted to black and white. The root network traits of the plants were then analyzed using the GiA Roots software (Galkovskyi et al., 2012). The further processing of the images encompassed their segmentation employing global thresholding (Binary_inverted) and Gaussian adaptive thresholding. For the comparative analysis, the total network area and total network length was used as readout. With respect to the biological variability of the plant root system, at least 24 individual plants per genotype and growth condition were analyzed.

### 2.5 Trypan blue staining

To confirm root colonization, 10-12 small root samples from control and infected plants were used. After thoroughly washing the root samples with deionized water, they were cut in 1 cm long pieces and incubated overnight in 10 N KOH. The explants were then rinsed 5 times with sterile H_2_O, before they were incubated for 5 min in 0.1 N HCl. Subsequently, the samples were incubated in a 0.05% Trypan blue solution (w/v), before they were partially decolorized with lactophenol over ten minutes. Before the specimen were mounted on glass slides and examined by microscopy, they were washed once with 100% ethanol and thrice with sterile H_2_O and stored in 60% glycerol (v/v).

### 2.6 Analysis of luciferase activity

For the *in vivo*-assessment of possible changes in auxin signaling activity in response to an infection with *S. indica*, bioluminescence measurements using the *DR5::Luc* reporter line were performed (Moreno-Risueno et al., 2010). To this end, the infection of *DR5::Luc* with the fungus was performed as previously described. In brief, at one, three, and six dpi, the luciferase activity was monitored both in the *S. indica*- and the mock-infected seedlings using a cooled CCD camera (NightOwl II LB 983 NC-100; Berthold Technologies). To visualize the luciferase activity, the plates were sprayed with 100 µM luciferin and imaged after an incubation time of 40 min. In the same manner, the effect of *S. indica*-infections of *pGH3.5::Luc* and *pGH3.17::Luc* reporter lines was carried out.

### 2.7 Confocal laser scanning microscopy

Differences in the local expression profiles of *GH3.5*, *GH3.17*, and *PIN2* in mock- and *S. indica*-infected roots were analyzed by using a Leica SP8 microscope with the Leica Application Suite (Las AF Lite) X software and the corresponding reporter lines *pGH3.5::3*×*YFP*, *pGH3.17::3*×*YFP*, and *pPIN2::PIN2-GFP*. On the one hand, the yellow fluorescent protein (YFP) was excited at 514 nm using an Argon multiline laser and detected using a 516-620 nm broadband filter. On the other hand, an excitation wavelength of 488 nm was used for the green fluorescent protein (GFP) and the detection of the emitted GFP fluorescence was achieved by employing a 494-596 nm broadband filter.

### 2.8 Mass spectrometric quantification of auxin and auxin conjugates

Free auxin levels were measured as previously described (Pérez-Alonso et al., 2021a). For this, 50 mg of plant material were harvested and shock-frozen in liquid nitrogen. After auxin extraction into methanol, one aliquot of the samples (60%) was spiked with 50 pmol of (^2^H_2_)-IAA as a stable isotope-labeled internal standard and analyzed by gas chromatography–tandem mass spectrometry (GC–MS/MS). A second aliquot (40%) was used to determine the amount of base-labile IAA-conjugates following a previously published procedure (Müller et al., 1998). It is generally inferred that the total IAA-conjugate amount in these fractions refers to the measure of the amount of IAA peptidyl-plus IAA glucosyl-conjugates. Other plant hormones and related compounds, including some plant hormone amino acid conjugates, were measured using an ultra-high performance-liquid chromatography (UHPLC)-electrospray interface quadrupole-orbitrap mass spectrometer (UHPLC/Q-Exactive; Thermo Scientific) setup equipped with an ODS column (ACQUITY UPLC HSS T3, 1.8 µm, 2.1 × 100 mm; Waters) as described previously (Kojima et al., 2009; Shinozaki et al., 2015).

### 2.9 RNA isolation and gene expression analysis by qRT-PCR

For each genotype and condition, 100 mg of plant tissue of either two or ten days-old sterilely grown seedlings were harvested for total RNA extraction as previously described (Oñate-Sánchez & Vicente-Carbajosa, 2008). First strand synthesis was conducted using M-MLV reverse transcriptase and oligo(dT)_15_ primer, following the instructions of the manufacturer (Promega). Two nanograms of cDNA were used as template in each qRT-PCR. cDNA amplification was performed using the FastStart SYBR Green Master solution (Roche Diagnostics) and a Lightcycler 480 Real-Time PCR system (Roche Diagnostics), according to the supplier’s instructions. The relative transcript quantification was calculated employing the comparative 2^-ΔΔ*C*T^ method (Livak & Schmittgen, 2001). As reference genes, we used *APT1* (At1g27450) and *GAPC2* (At1g13440) (Czechowski et al., 2005; Jost et al., 2007). The quantitative gene expression analysis was carried out as previously described (Pérez-Alonso, et al., 2021b), using three biological replicates. In addition, three technical replicates per biological replicate were analyzed. See **Supporting Information:Data Sheet 1** for primer sequences.

### 2.10 RNA-seq analysis

In this study, we performed genome-wide expression studies employing mRNA sequencing (RNA-seq). To do so, total RNA from ten days-old mock- and *S. indica*-infected wild-type and YUC9ox seedlings were extracted as described above and quantified using a Nanodrop ND-1000^®^ UV/Vis spectrophotometer (ThermoFisher). RNA quality was additionally checked on a Bioanalyzer 2100 (Agilent) by the Novogene Genomics Service (Cambridge, UK). Library construction and sequencing (150-nt paired-end reads) on Illumina NovaSeq™ 6000 platforms was subsequently performed by the Novogene Genomics Service, that also provided basic data analysis applying their RNA-seq pipeline.

To analyze overlapping pattern in differentially expressed genes, Venn plots have been generated using the Venn (http://bioinformatics.psb.ugent.be/webtools/Venn/) online tool. For the gene ontology (GO) enrichment analysis we used either the g:Profiler online tool (Raudvere et al., 2019) or the Metascape gene annotation and analysis resource (Zhou et al., 2019). GO chord plots were generated using the SRplot graphical interphase (https://www.bioinformatics.com.cn/en), while the principal component analysis and the dot plots were generated using the GraphBio application (Zhao & Wang, 2022).

### 2.11 Statistical analysis

The statistical assessment of the data was performed using the JASP v0.16.1 software (https://jasp-stats.org/). Student’s *t*-test was employed to compare two means. Results were considered significant when the *p*-value < 0.05.

## 3. Results

### 3.1 *S. indica* induced root growth does not depend on the induction of auxin biosynthesis-related genes

The growth promoting effect of the beneficial root colonizing fungus *S. indica* is well documented, including considerable stimulation of plant root growth (Pérez-Alonso et al., 2020; Su et al., 2017). Furthermore, the crucial role of auxin in controlling root development is widely accepted (Roychoudhry & Kepinski, 2022). For these reasons, it was tempting to speculate that the promotion of root growth triggered by the fungus is achieved by inducing auxin biosynthesis in the host plant. To address the question of possible fungus-mediated transcriptional activation of auxin biosynthesis-related genes in Arabidopsis seedlings, we performed a series of RNA-seq experiments two and ten days after infection (dpi) (Pérez-Alonso et al., 2022). However, neither the global analysis of gene set enrichment (Gene Ontology (GO) analysis) nor the directed analysis of 31 auxin biosynthesis-related genes in this data set provided evidence for substantial induction of these genes (**Figure 1, Supporting Information:Image 1**).

**Figure 1.**
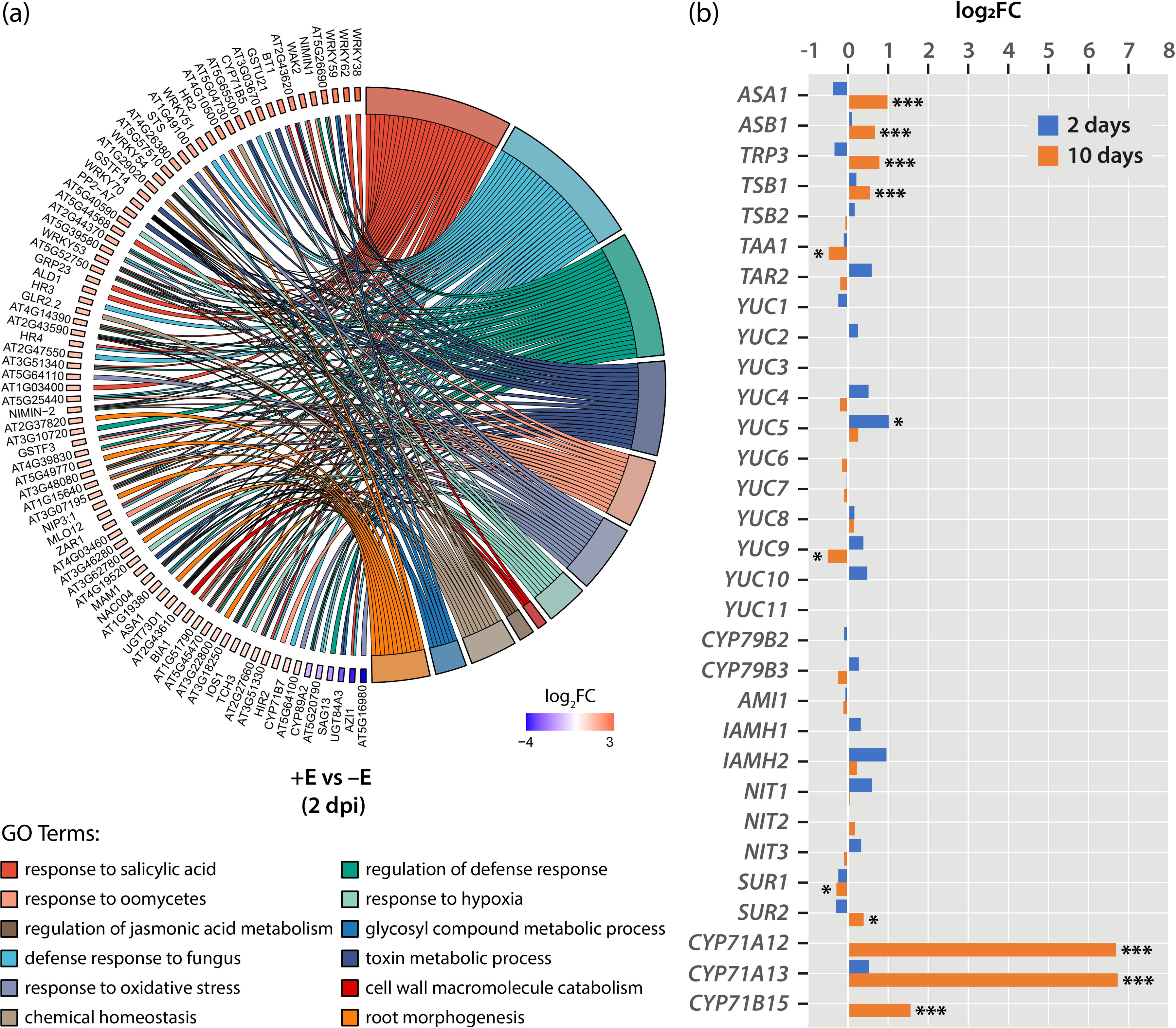
Transcriptional analysis of *S. indica*-infected Arabidopsis seedlings. (**a**) Chord plot showing the most enriched biological processes (GO terms) with their DEGs in *S. indica*-infected versus control plants (2 dpi). In each chord, enriched GO biological process terms are shown on the right, and the DEGs contributing to this enrichment are shown on the left. On the left side, each DEG is represented by a rectangle which color is correlated to the value of the differential expression (log_2_FC). *S. indica* up-regulated genes are displayed in red whereas down-regulated genes are displayed in blue. Chords connect gene names with biological process GO term groups. Each GO term is represented by one colored line. (**b**) Expression levels of 31 auxin biosynthesis-related genes in *S. indica*-infected seedlings relative to control plants at 2 (blue) and 10 dpi (orange). The bars show means of n = 3 independent measurements. Asterisks mark the genes with a significantly altered expression. Student’s *t*-test: *p ≤ 0.05, ***p ≤ 0.001.

The co-colonization of Arabidopsis seedlings with *S. indica* does not result in the enrichment of DEGs associated with auxin metabolism-related GO classifications (**Figure 1a**). At two dpi, only significant induction of the *YUCCA5* (*YUC5*) gene was registered (**Figure 1b**). At the later time point (10 dpi), the induction of the genes coding for the α- and β-subunit of *ANTHRANILATE SYNTHASE* (*ASA1* and *ASB1*) and the α- and β-chains of *TRYPTOPHAN SYNTHASE* (*TRP3* and *TSB1*) suggested an increased formation of L-Trp. However, the metabolic flux appears to be directed toward the formation of defense-related compounds, including camalexin and glucosinolates, because the relevant cytochrome P450 genes *SUR2*, *CYP71A12*, *CYP71A13*, and *CYP71B15* appeared to be substantially induced (Barlier et al., 2000; Böttcher et al., 2009; Müller et al., 2019), while the genes *TRYPTOPHAN AMINOTRANSFERASE OF ARABIDOPSIS 1* (*TAA1*) (Stepanova et al., 2008) and *YUC9* (Hentrich et al., 2013), involved in *de novo* auxin biosynthesis, appeared to be slightly repressed, thus confirming previously published results (Fröschel et al., 2021; Lahrmann et al., 2015).

### 3.2 Auxin conjugation and signaling is altered in *S. indica* induced seedlings

To gain further insight into auxin dynamics during the establishment of the symbiosis between *S. indica* and Arabidopsis, free IAA as well as total IAA conjugate levels were quantified by tandem mass spectrometry (**Figure 2a, Supporting Information:Image 2**). Although no major induction of auxin biosynthesis-related genes was observed, free IAA levels were significantly elevated in Arabidopsis seedlings co-cultivated with *S. indica* after 3 days, before the IAA content dropped, showing no longer significant differences between *S. indica*- and mock-infected plants. The analysis of total conjugated IAA provided no evidence for significant differences between the two conditions tested. However, when directly analyzing the contents of indole-3-acetyl-L-aspartic acid (IAA-Asp), a significant increase in the compound was recorded in later stages of infection, suggesting a possible involvement of auxin conjugation in the establishment of symbiosis. To support the notion of altered auxin contents during the infection, auxin signaling was investigated using a *DR5::Luc* reporter line (Moreno-Risueno et al., 2010). As demonstrated in **Figure 2b,c**, auxin signaling appeared to be induced at 1 dpi, consistent with the observed increased IAA contents. At 3 dpi, the quantification of auxin signaling activities showed no significant differences between infected and non-infected roots. However, at 6 dpi, auxin signaling was found to be again significantly stronger in the *S. indica*-infected plants.

**Figure 2.**
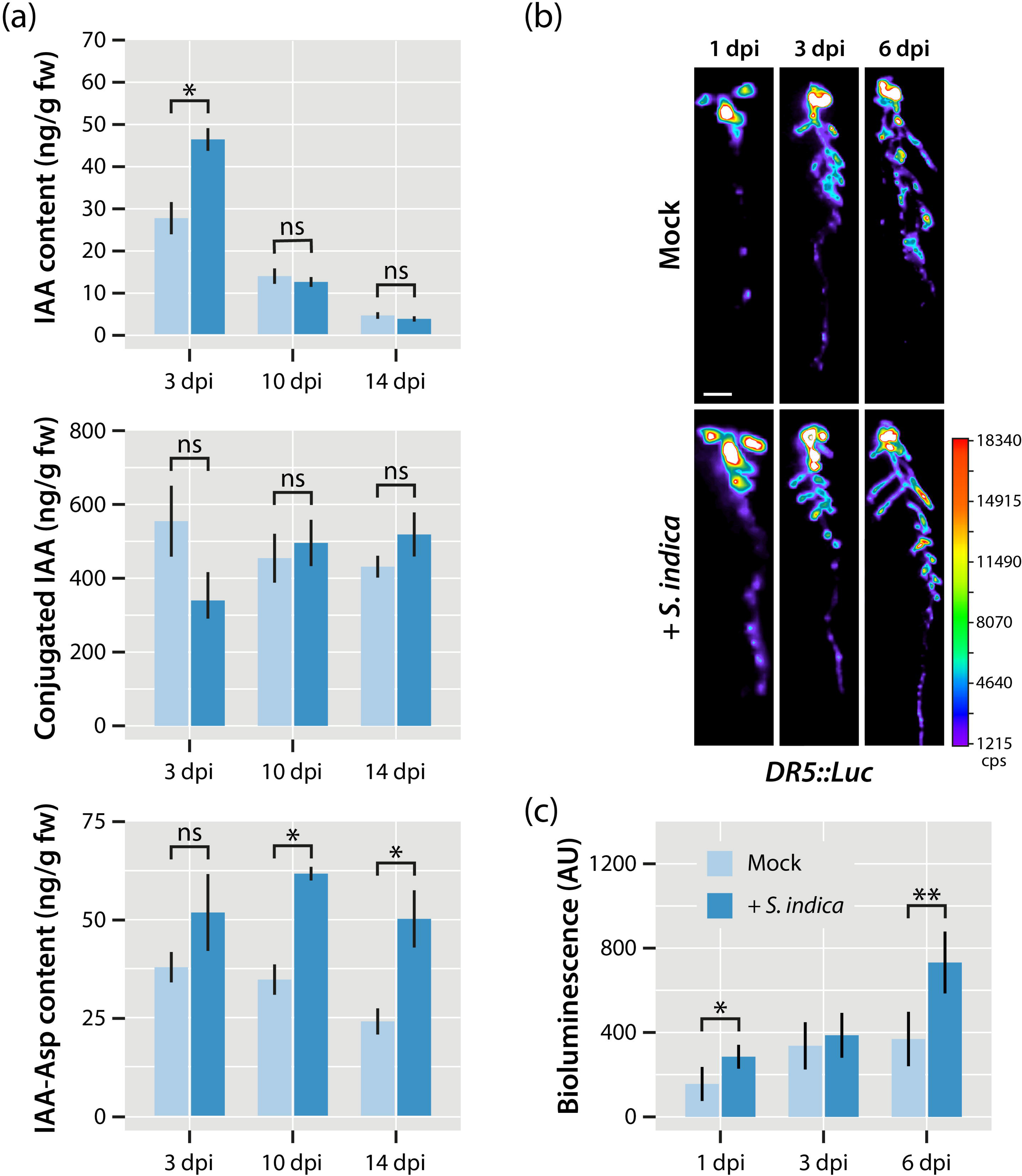
Auxin contents and auxin signaling in *S. indica*-infected Arabidopsis seedlings. (**a**) Mass spectrometric assessment of free auxin (top panel), total conjugated auxin (middle panel), and indole-3-acetyl-l-aspartic acid (IAA-Asp) (bottom panel). The bars show means of n = 3 independent measurements. Asterisks mark the conditions with significantly altered compound levels. Student’s *t*-test: *p ≤ 0.05. (**b**) The images show representative *DR5::Luc* bioluminescence values obtained after long-term imaging (5 min exposure). Scale bar = 1 cm. Quantification of *DR5::Luc* signals in the root systems of *S. indica*- and mock-infected Arabidopsis seedlings (n = 5). AU = Arbitrary Units.

These results confirm previous studies that reported an auxin signaling peak within the first 24 h after infection (Meents et al., 2019) and provided evidence for the production of IAA by *S. indica* (Hilbert et al., 2012). Intriguingly, the auxin signaling activity was found to be significantly enhanced in later stages of the plant–fungus interaction, although auxin levels were observed to drop and show no difference in mock- and fungus-infected seedlings.

### 3.4 Co-cultivation with *S. indica* can restore a wild type-like phenotype in high auxin mutants

A previous work provided evidence that the co-cultivation of *sur1-1*, a mutant with high auxin contents, with *S. indica* can rescue the strong auxin phenotype (Vadassery et al., 2008). However, given that the *sur1-1* mutant also interferes with the defense response of Arabidopsis, as mutant plants are impaired in the production of indole glucosinolates, an alternative auxin-overproducing mutant, YUC9ox, was used to address the question of whether infection with *S. indica* is indeed sufficient to restore a wild-type-like phenotype in this high auxin line (Hentrich et al., 2013). As shown in **Figure 3a**, infection of YUC9ox seedlings with *S. indica* resulted in a considerable change in the architecture of the root system, suggesting the restoration of normal auxin contents in YUC9ox. In order to investigate the growth regulating effect of *S. indica* in high auxin mutant seedlings, the transcriptional changes involved were investigated by comparing mock- and *S. indica*-infected YUC9ox seedlings with similarly treated wild-type Arabidopsis seedlings at 10 dpi by RNA-seq. A principal component analysis of the normalized expression units obtained (FPKM) for the different RNA-seq reactions provided evidence for a clear separation of four distinct groups, highlighting differences both at the genetic level (PC1, YUC9ox versus Col-0) and at the treatment level (PC2, *S. indica-* versus mock-infected seedlings) (**Figure 3b**). When focusing on differentially expressed genes (DEGs) under the given conditions (false discovery rate (FDR) ≤ 0.01; log_2_(fold change) ≥ |1.25|), it became apparent that the effect of *S. indica* on YUC9ox and wild-type control plants (Col-0) was relatively low, triggering differential expression of only a small number of genes with 352 and 189 induced genes and 10 and 34 repressed genes, respectively, for *S. indica-* versus mock-infected YUC9ox and Col-0 seedlings. However, contrasting DEGs between genotypes (YUC9ox versus Col-0) and growth conditions (+endophyte (E) vs –E) elevated the number of identified DEGs to 671 and 704 induced genes and 451 and 750 repressed genes, respectively, for the comparison between mock- and *S. indica*-infected seedlings (**Figure 3c**). The detailed analysis of the latter comparison is shown in the Venn plot in **Figure 3d**. Here, the 402 DEGs that were induced under mock- and *S. indica*-infected conditions attracted our special interest, as this group is supposed to comprise genes that respond to both high auxin levels and the co-cultivation with the fungus. The GO analysis of those genes revealed the overrepresentation of DEGs related to the response to auxin (**Figure 3e**).

**Figure 3.**
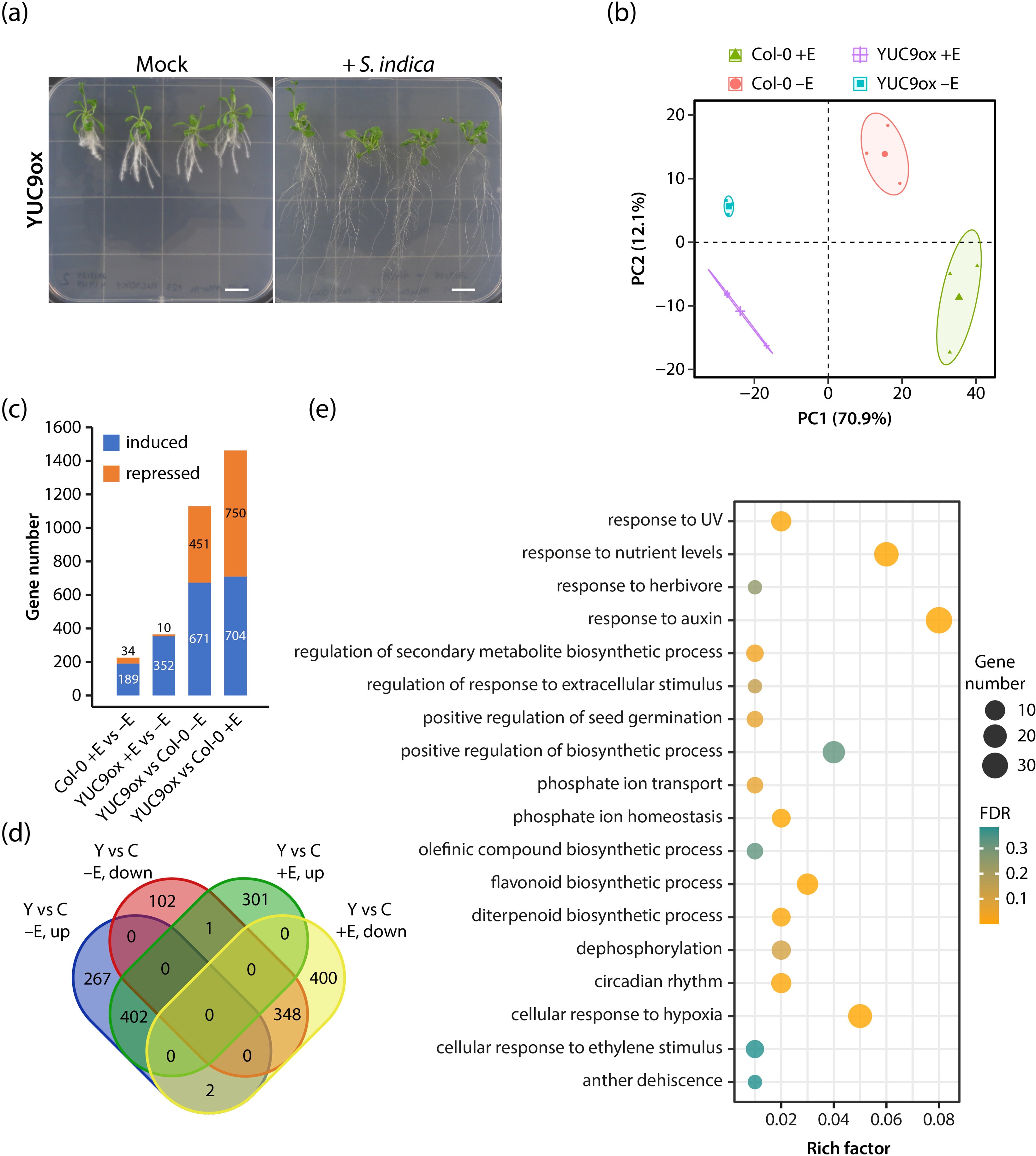
Analysis of the effects of *S. indica* infection on YUC9ox and wild-type Arabidopsis seedlings. (**a**) Phenotype of auxin overproducing YUC9ox seedlings that were either mock-infected or inoculated with 20 µl of a solution containing 2 × 10^5^ *S. indica* spores ml^-1^ at 10 dpi. The scale bars = 1 cm. (**b**) Principal component analysis (PCA) of transcriptome-wide normalized gene expression counts assessing the difference in mock (–E) and *S. indica* (+E) infected YUC9ox and wild-type Col-0 seedlings. Each dot represents a sample, color and symbol are coded by genotype and condition (Circle: wild-type Arabidopsis without endophyte (Col-0 –E); Triangle: wild-type Arabidopsis with endophyte (Col-0 +E); Square: YUC9ox without endophyte (YUC9 –E); Cross: Square: YUC9ox with endophyte (YUC9ox +E)). The x-axis denotes the PC1; y-axis denotes values for PC2. (**c**) DEGs statistics for the YUC9ox mutant and Col-0 control plants after co-cultivation with *S. indica* for 10 days. (c) Venn diagram showing the numbers of DEGs in YUC9ox (Y) versus Col-0 (C) plants that were either mock (–E) or *S. indica* (+E) infected at 10 dpi. Induced (up) and repressed (down) DEGs were separately evaluated. (**e**) GO enrichment analysis of the 402 induced DEGs in mock- and *S. indica*-infected YUC9ox seedlings versus similarly treated Col-0 control plants. Each circle in the figure represents a distinct GO term, and the circle size indicates the number of genes enriched in the corresponding GO term. The significance of the observed gene enrichment is represented by a color gradient referring to the FDR (q-value).

Among the 33 DEGs related to the ‘response to auxin’ GO term classification, 8 small auxin up-regulated genes (*SAUR24*, *SAUR29*, *SAUR34*, *SAUR35*, *SAUR66*, *SAUR68*, *SAUR69*, *SAUR76*) were found, along with 5 Aux/IAA repressor genes (*IAA1*, *IAA5*, *IAA6*, *IAA19*, *IAA29*) and 5 *GH3* genes (*GH3.1*, *GH3.3*, *GH3.5*, *GH3.7*, *GH3.8*). Of all the candidates identified, only GH3 enzymes are known to directly act on free auxin and conjugate the active hormone with amino acids, thus inactivating it. Taken together, our results suggested that GH3 acyl amido transferases are likely to play an important role in controlling endogenous auxin levels in *Arabidopsis* seedlings infected with *S. indica*.

### 3.5 Impact of selected *GH3* genes on *S. indica*-mediated growth promotion

To further evaluate a possible role of *GH3* genes in the root growth promotion mediated by *S. indica*, we identified nine *GH3* genes, i.e., *GH3.1*, *GH3.2*, *GH3.3*, *GH3.4*, *GH3.5*, *GH3.6*, *GH3.9*, *GH3.11*, and *GH3.17*, in the RNA-seq data of fungus infected Col-0 plants that appeared to be induced by the plant–microbe interaction. Consistency of the RNA-seq data was confirmed by the qRT-PCR analysis of the differential expression of the genes in shoots and roots of mock- and *S. indica*-infected wild-type plants at 2 and 10 dpi. Hierarchical clustering of the normalized differential expression data revealed the existence of four distinct groups (**Figure 4**). While *GH3.1* and *GH3.6* appear to be mainly expressed in shoots, all other tested genes displayed a more pronounced expression in root tissues. Among those *GH3* genes, *GH3.2*, *GH3.11*, and *GH3.12* are more strongly expressed at later stages, whereas *GH3.4*, *GH3.5*, *GH3.17*, *GH3.3*, and *GH3.9* are induced at 2 dpi and display a reduction in their expression levels at 10 dpi. Apart from that, the two latter genes, *GH3.3* and *GH3.9*, differ from the other ones because they seem to be substantially repressed in shoots at 2 dpi.

**Figure 4.**
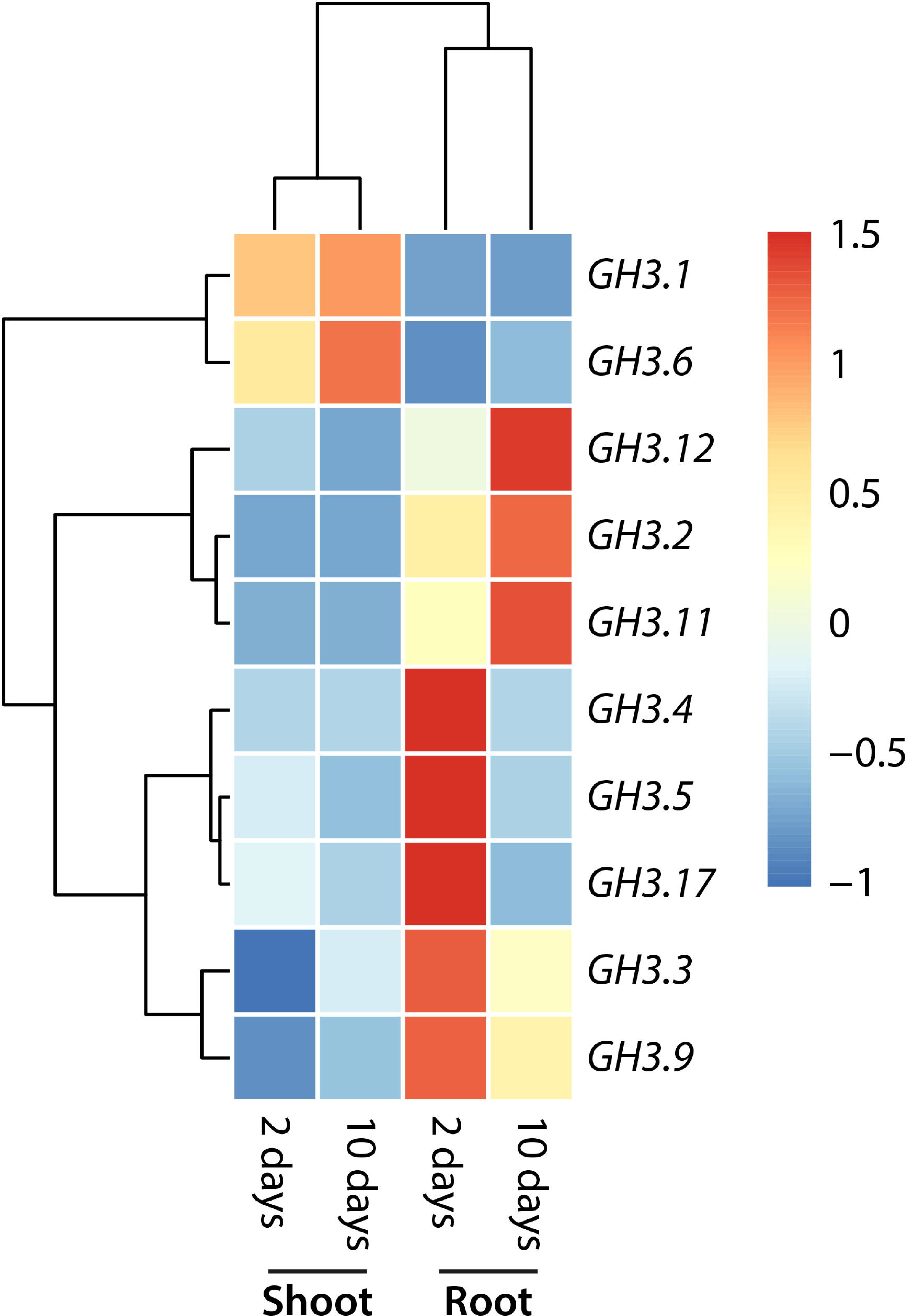
Hierarchical clustering of normalized expression values at 2- and 10-dpi in root and shoot tissues of *S. indica*-versus mock-infected Col-0 seedlings. The heatmaps reflect gene expression values normalized to the mean across all time points (day 2 and 10 in roots and shoots) for genes that met the cutoff in at least one time point (p ≤ 0.05 and fold change ≥ 2).

Especially the three *GH3* genes *GH3.4*, *GH3.5* and *GH3.17* sparked our interest, as they quickly responded to the infection and have already been reported to accept IAA as substrate or to be involved in root elongation (Guo et al., 2022; Staswick et al., 2005; Staswick et al., 2002). To investigate the involvement of GH3.4, GH3.5 and GH3.17 in the establishment of the plant–fungus interaction in more detail, we analyzed the promotion of fungus-mediated root growth in the corresponding *gh3.4*, *gh3.5*, and *gh3.17* knockout mutants. *S. indica-* infected *gh3.4* and wild-type plants showed no obvious difference. In both cases, the fungus triggered significant root growth of the seedlings (**Figure 5, Supporting Information:Image 3**).

**Figure 5.**
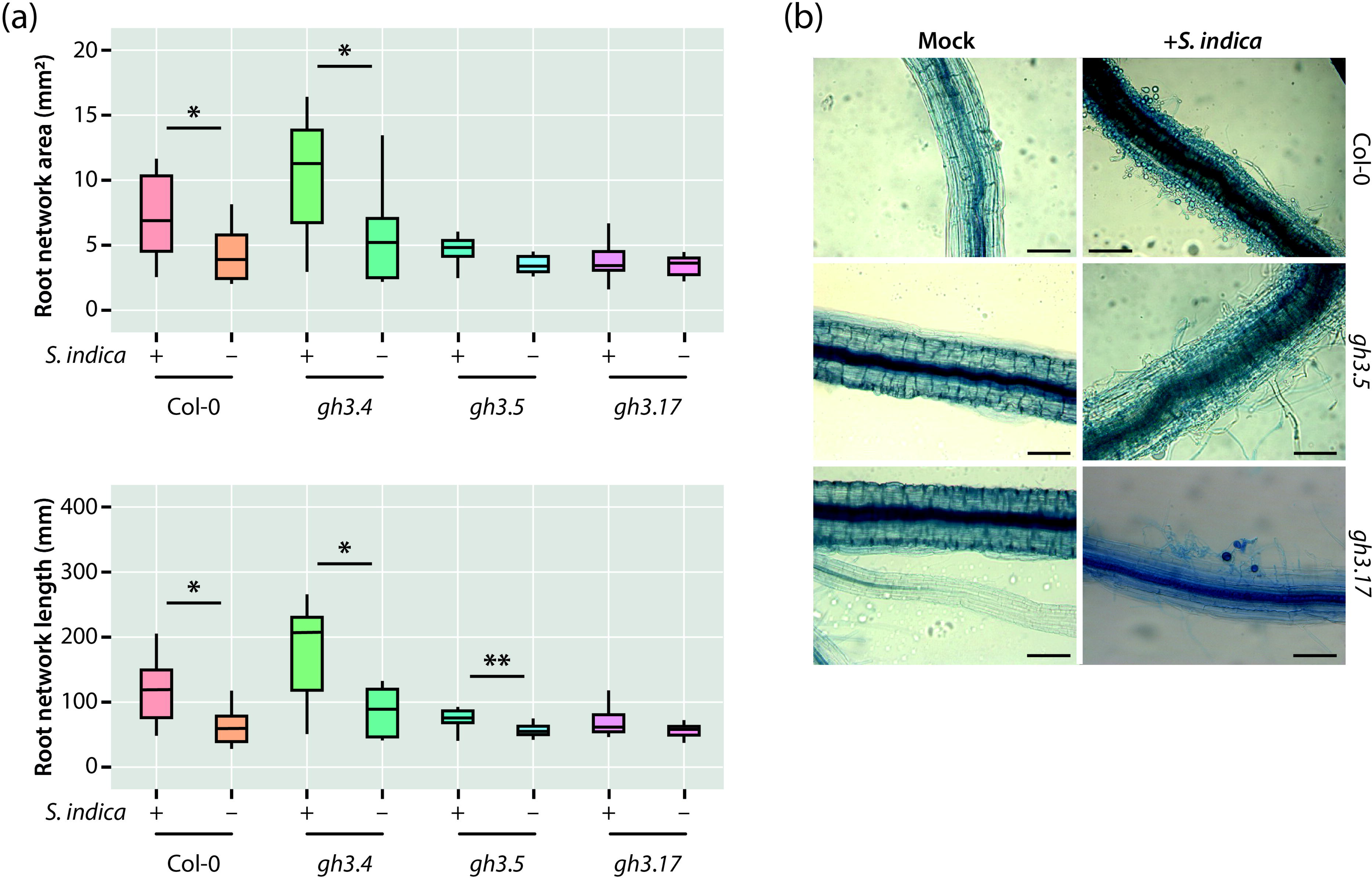
Analysis of the colonization of Col-0 and *gh3* mutant roots with *S. indica*. (**a**) Total root network area and length of Arabidopsis wild-type and *gh3* knockout mutant plants after 10 days of co-cultivation with *S. indica* or mock treatment. The box plots show the median, quartiles, and extremes of the compared data sets (n = 24). Asterisks indicate significant differences between *S. indica*- and mock-treated samples. Student’s *t*-test: *p ≤ 0.05, **p ≤ 0.01. (**b**) Trypan blue stain of root segments at 1−2 cm distance from the root tip. The figure shows representative pictures for the three studied genotypes after 10 days of co-cultivation with *S. indica*. Scale bars = 1 cm.

As evidenced in **Figure 5a**, the growth promoting effect of the root endophyte was largely missing in the *gh3.5* and *gh3.17* mutants. The *gh3.5* mutant plants still showed the tendency to respond positively to fungus infection, showing a significantly increased total root length after infection, although the values remained far behind those observed in wild-type control and *gh3.4* seedlings. However, mock- and *S. indica*-infected *gh3.17* plants were indistinguishable, showing no significant growth response. It must be noted that the missing response to *S. indica* cannot be attributed to impaired colonization of the roots of *gh3.5* and *gh3.17*. The microscopic inspection of Trypan blue stained root fragments provided indisputable evidence for the colonization of mutant roots (**Figure 5b**). Based on these results, it must be concluded that the two GH3 enzymes, GH3.5 and GH3.17, play an important role in establishing the mutual interaction between *S. indica* and Arabidopsis.

### 3.6 Fungus-induced auxin conjugation is necessary for plant growth promotion

Our results suggested the involvement of auxin conjugation in the development of the mutual interaction between *S. indica* and Arabidopsis. To challenge this hypothesis, we decided to explore the root growth promoting effect of the fungus on Arabidopsis seedlings that constitutively overexpress *IAR3*. The *IAR3* gene encodes an auxin conjugate hydrolase that is normally expressed in Arabidopsis roots, where it hydrolyzes indole-3-acetyl-L-alanine (IAA-Ala) to release free IAA (Davies et al., 1999). The genetically engineered increased release of IAA in *35S::IAR3* lines was supposed to counteract the induced conjugation of free IAA after root colonization with the fungus. As displayed in **Figure 6**, we used two independent *35S::IAR3* overexpression lines with different levels of transgene expression. While line 6.2 showed only moderate expression of the *IAR3* cDNA, line 1.3 displayed a strong overexpression of the transgene (**Figure 6b**).

**Figure 6.**
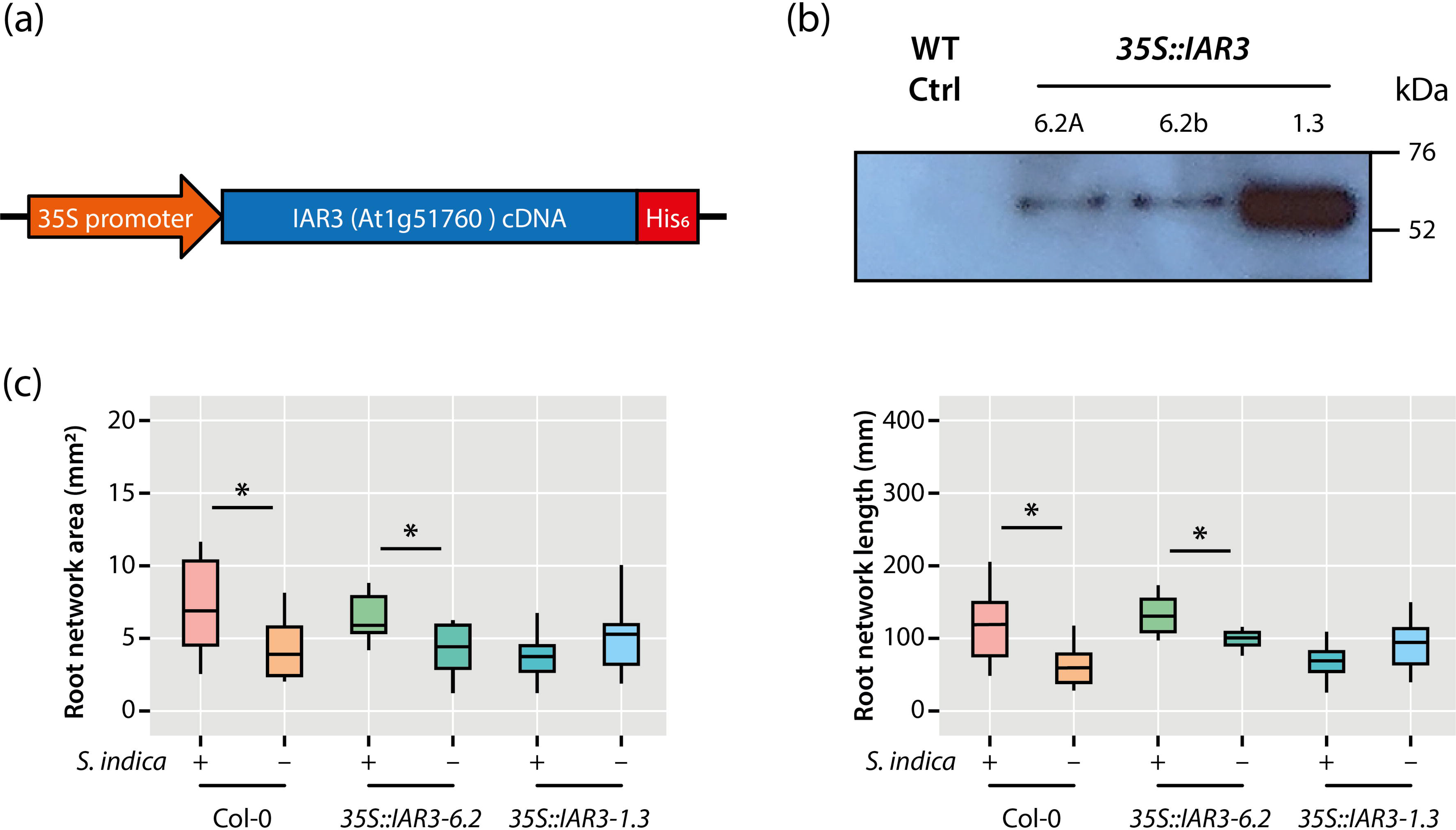
Analysis of the *S. indica*-mediated root growth promotion of Arabidopsis Col-0 and *35S::IAR3* roots. (**a**) Topology of the generated *IAR3* transgene construct. (**b**) Protein immunodetection of the IAR3-His_6_ transgene in wild-type and two independent *35S::IAR3* lines using an anti-His antibody. (**c**) Total root network area and length of Arabidopsis wild-type and constitutively IAR3 overexpressing mutant lines after 10 days of co-cultivation with *S. indica* or mock treatment. The box plots show the median, quartiles, and extremes of the compared data sets (n = 24). Asterisks indicate significant differences between *S. indica*- and mock-treated samples. Student’s *t*-test: *p ≤ 0.05.

Consistent with the expression levels of the employed lines, we found an approximately 40-50% reduced root growth promotion in *S. indica*-infected seedlings of line 6.2, while the strong overexpressor line 1.3 showed no growth promotion at all, but rather a negative effect on root growth, when compared to wild-type plants. The altered growth promotion effects in the *IAR3* overexpressing lines suggest that the intimate control of auxin conjugation during the establishment of the interaction between Arabidopsis and *S. indica* is a relevant determinant for the development of the beneficial effect of the fungus on root growth

### 3.7 *S. indica* triggers the local induction of *GH3.5* and *GH3.17*

Next, we addressed the question of whether the observed induction of the *GH3.5* and *GH3.17* genes is locally restricted or if the induction of gene expression spreads over the entire root. To this end, we monitored the transcriptional response of the two genes using corresponding transgenic promoter-reporter lines (Pierdonati et al., 2019) over the course of the infection with *S. indica*. As displayed in **Figure 7**, the two genes show distinct locally restricted expression patterns at the primary root and lateral root tips, with *GH3.5* generally showing stronger expression levels under control conditions.

**Figure 7.**
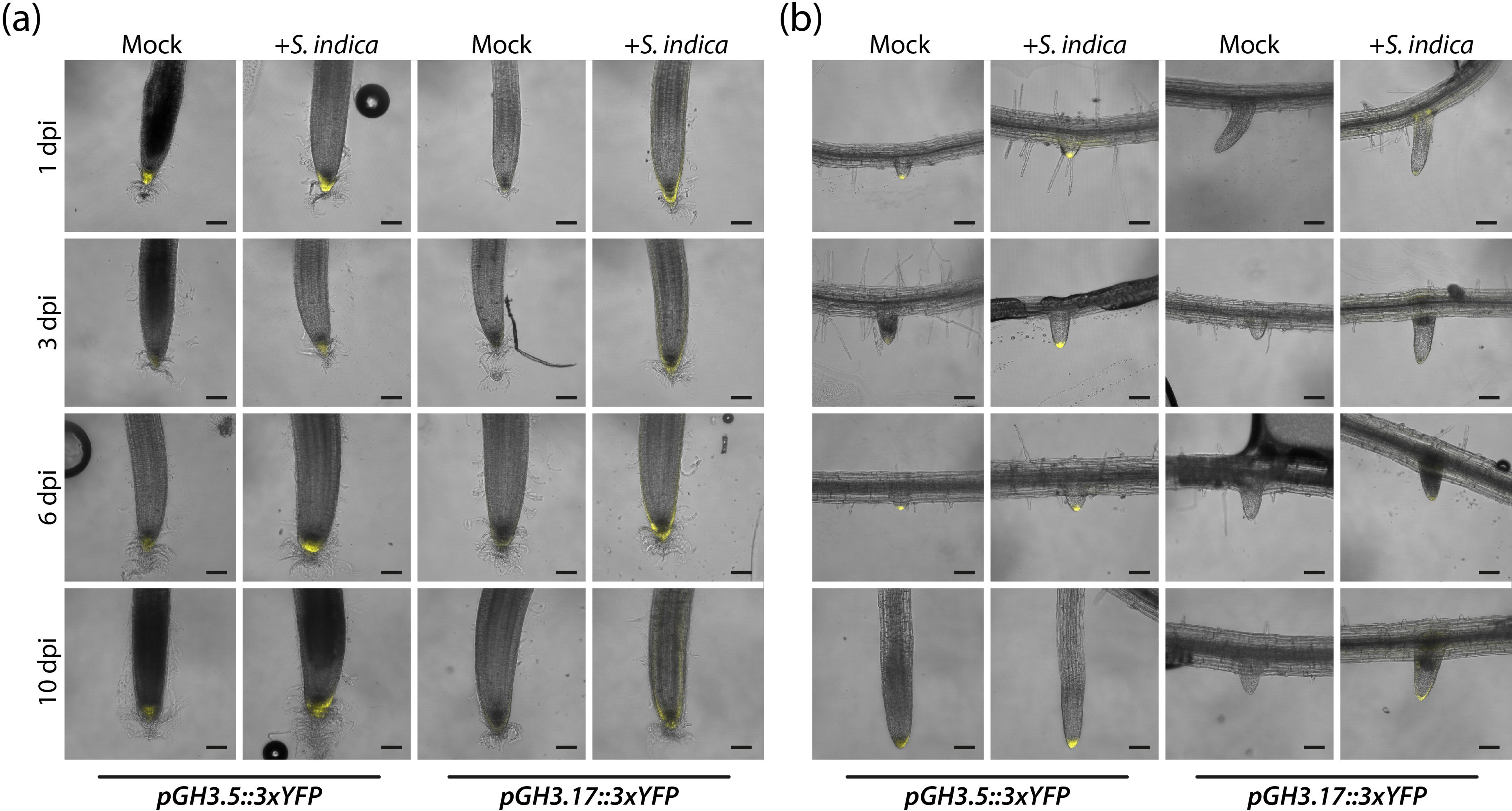
*S. indica* infection induces the expression of *GH3.5* and *GH3.17* in primary and lateral root. (**a**) Overlay of confocal and bright field images of primary root tips of mock- and *S. indica*-infected *pGH3.5::3*×*YFP* and *pGH3.17::3*×*YFP* promoter-reporter lines 1, 3, 6 and 10 dpi. (**b**) Overlay of confocal and bright field images of lateral root tips of mock- and *S. indica*-infected *pGH3.5::3*×*YFP* and *pGH3.17::3*×*YFP* constructs at 1, 3, 6 and 10 dpi. Scale bars = 100 µm.

Confirming our transcriptomics analyzes, we were able to detect the induction of the expression of both genes. In the primary root (**Figure 7a**), the expression of *GH3.5* was restricted mainly to columella cells, and fungus infection moderately increased the expression strength. In some cases, that is, at 1 and 10 dpi, the expression of *GH3.5* appeared to also spread to lateral root cap cells. *GH3.17* expression was restricted to an external single cell layer in the lateral root cap under control conditions. However, after infection with *S. indica,* the expression of *GH3.17* was much stronger induced than the expression of *GH3.5* and, in later stages (10 dpi), appeared to extend into the root cortex. With respect to the lateral root tips (**Figure 7b**), the expression of both genes was localized to the lateral root tip. However, it should be noted that the expression level of *GH3.17* at the lateral root tips was extremely low and difficult to detect. As with the primary roots, the effect of fungus infection on *GH3.17* expression was much stronger compared to the response of *GH3.5*. Nonetheless, it needs to be remarked that both the expression of *GH3.5* and *GH3.17* was also induced at the base of the emerging lateral root and the surrounding primary root cells. For *GH3.5*, on the one hand, this effect was only detectable at 1 dpi, while for *GH3.17*, on the other hand, this pattern was found throughout the experiment (**Supporting Information:Image 4**). The generated *pGH3.5::Luc* and *pGH3.17::Luc* lines confirmed our finding of the induction of the expression of the two genes in the root system. Intriguingly, the quantification of the bioluminescence in the complete root system of mock- and *S. indica*-treated promoter-reporter plants revealed a significant and sustained activation of the promoters at later stages of the infection. While the *pGH3.17::Luc* constructs start to show differences 6 dpi, the *pGH3.5::Luc* lines already exhibit a significantly stronger activation after 3 dpi, when compares to corresponding mock controls (**Supporting Information:Image 5**). These results unequivocally confirm that the effect of *S. indica* on the expression of *GH3.5* and *GH3.17* is spatially restricted and sustained over a longer period of time throughout the infection of the root.

### 3.8 *S. indica*-infection reduces the amount of *PIN2* in the root

The transcriptomics and life cell imaging experiments allowed us to identify an increased expression of *GH3.5* and *GH3.17* in *S. indica*-infected root tips. However, previous studies reported that *S. indica* does not invade the meristematic zone of root tips, unless developmentally programmed cell death is impaired (Charura et al., 2023; Jacobs et al., 2011). So, we asked the question how the induction of *GH3.5* and *GH3.17* expression in the root tips could be triggered. Considering the already described responsiveness of *GH3* genes towards auxin treatments (Hagen & Guilfoyle, 2002), we speculated that the local induction of the two *GH3* genes is presumably caused by the formation of local auxin accumulations at the tips of the roots. This could be provoked by an increase in acropetal auxin transport toward the tip through induction of *PIN1* or a reduction in basipetal auxin redirection through a repression of *PIN2* (Michniewicz et al., 2007). To test this hypothesis, we returned to our RNA-seq data set (Pérez-Alonso et al., 2022) and checked the expression levels of the five *PIN* genes, *PIN1*, *2*, *3*, *4*, and *7*, involved in intercellular polar auxin transport. As shown in **Figure 8a**, the RNA-seq data displayed only moderate but significant suppression for *PIN2*, while the differential expression of the other *PINs* showed no significant alteration in response to the infection with the fungus. Subsequent analysis of the expression by qRT-PCR confirmed the repression of *PIN2* and pointed to a possibly very weak suppression of *PIN1* in response to the *S. indica* infection.

**Figure 8.**
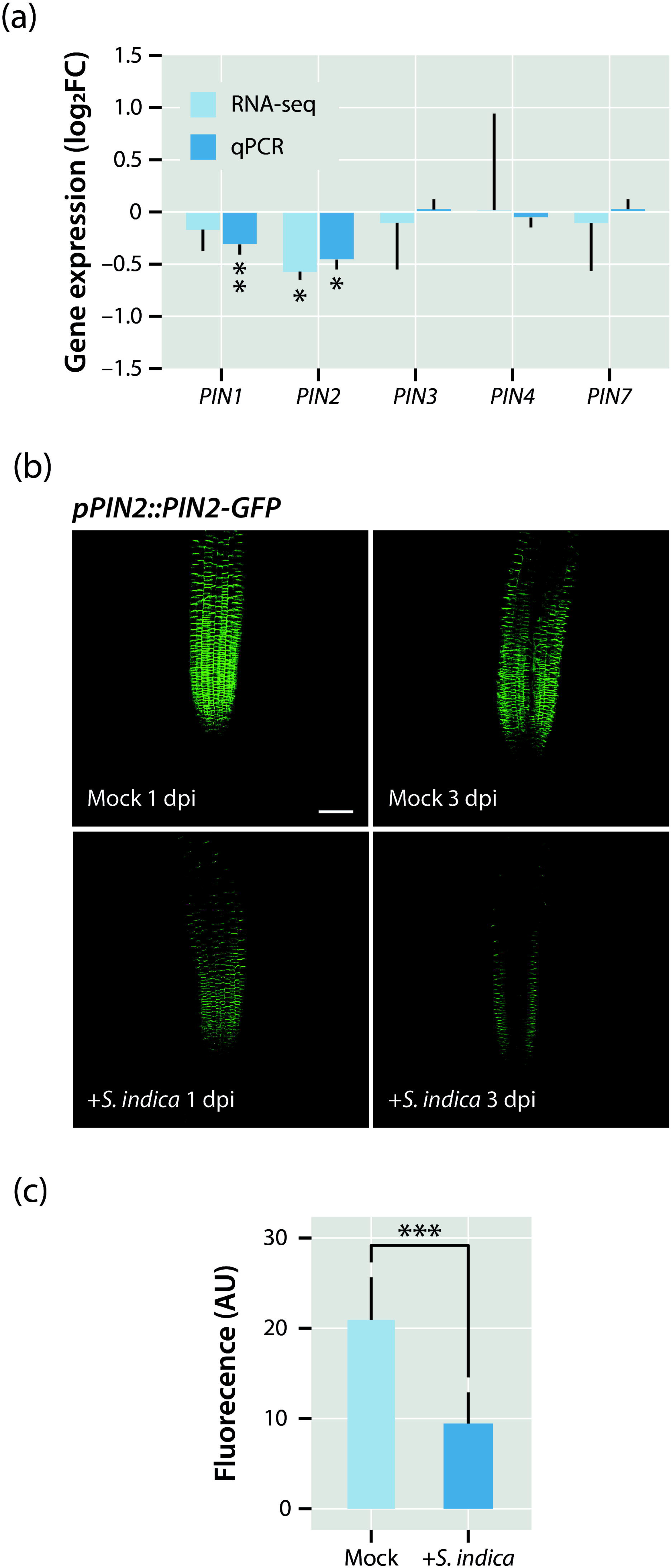
Qualitative and quantitative analysis of *PIN* transporter responses to an infection with *S. indica*. (**a**) Analysis of transcriptional alterations of selected *PIN* genes due to an infection with *S. indica* by RNA-seq and qRT-PCR. The bars show means of n = 3 independent measurements. Asterisks mark the genes with a significantly altered expression. Student’s *t*-test: *p ≤ 0.05, **p ≤ 0.01. (**b**) Confocal laser scanning microscopy images of primary root tips of mock- and *S. indica*-infected *pPIN2::PIN2-GFP* promoter-reporter plants 1 and 3 dpi. Scale bar = 100 µm. (**c**) Quantification of GFP fluorescence in root tips of mock- and *S. indica*- infected *pPIN2::PIN2-GFP* plants at 3 dpi. The bars show means of n = 5 independent measurements. Asterisks mark significantly altered fluorescence levels. Student’s *t*-test: ***p ≤ 0.001.

To provide further experimental proof for the transcriptional response of *PIN2* to *S. indica* at a cellular resolution, we examined the expression of a corresponding *pPIN2::PIN2-GFP* reporter line under mock control conditions and in fungus-infected plants. The fluorescent signal of the *PIN2* reporter at the root tips appeared considerably reduced when the plants were infected with the fungus (**Figure 8b,c**), thereby confirming our transcriptomics data and hinting at an important role of PIN2 in the formation of local auxin maxima in the root tips which, in consequence, can lead to the induction of *GH3* gene expression in these regions.

### 3.9 Partial loss of PIN2 activity is sufficient to trigger *GH3.5* and *GH3.17* expression and results in increased plant biomass

Finally, we aimed at testing whether the alteration of the auxin distribution in the root tips through the partial loss of PIN2 is sufficient to induce *GH3.5* and *GH3.17* expression. At the same time, our goal was to investigate if these changes can also affect the biomass of the seedlings, because a previous study pointed out that the accumulation of auxin in the root tips promotes growth in transgenic maize (Li et al., 2018). For this, we quantified *GH3.5* and *GH3.17* expression in the two PIN2 mutants *agr1-1* and *agr1-2* (Bell & Maher, 1990; Chen et al., 1998) and measured their fresh and dry weight in comparison to the corresponding Arabidopsis wild-type, Ler-0 (**Figure 9**). While *agr1-2* is characterized by a premature stop codon after 87 nucleotides, *agr1-1* is a less severe mutant allele that contains a functional G519D mutation in the seventh of the ten α-helical domains of the transporter (Utsuno et al., 1998).

**Figure 9.**
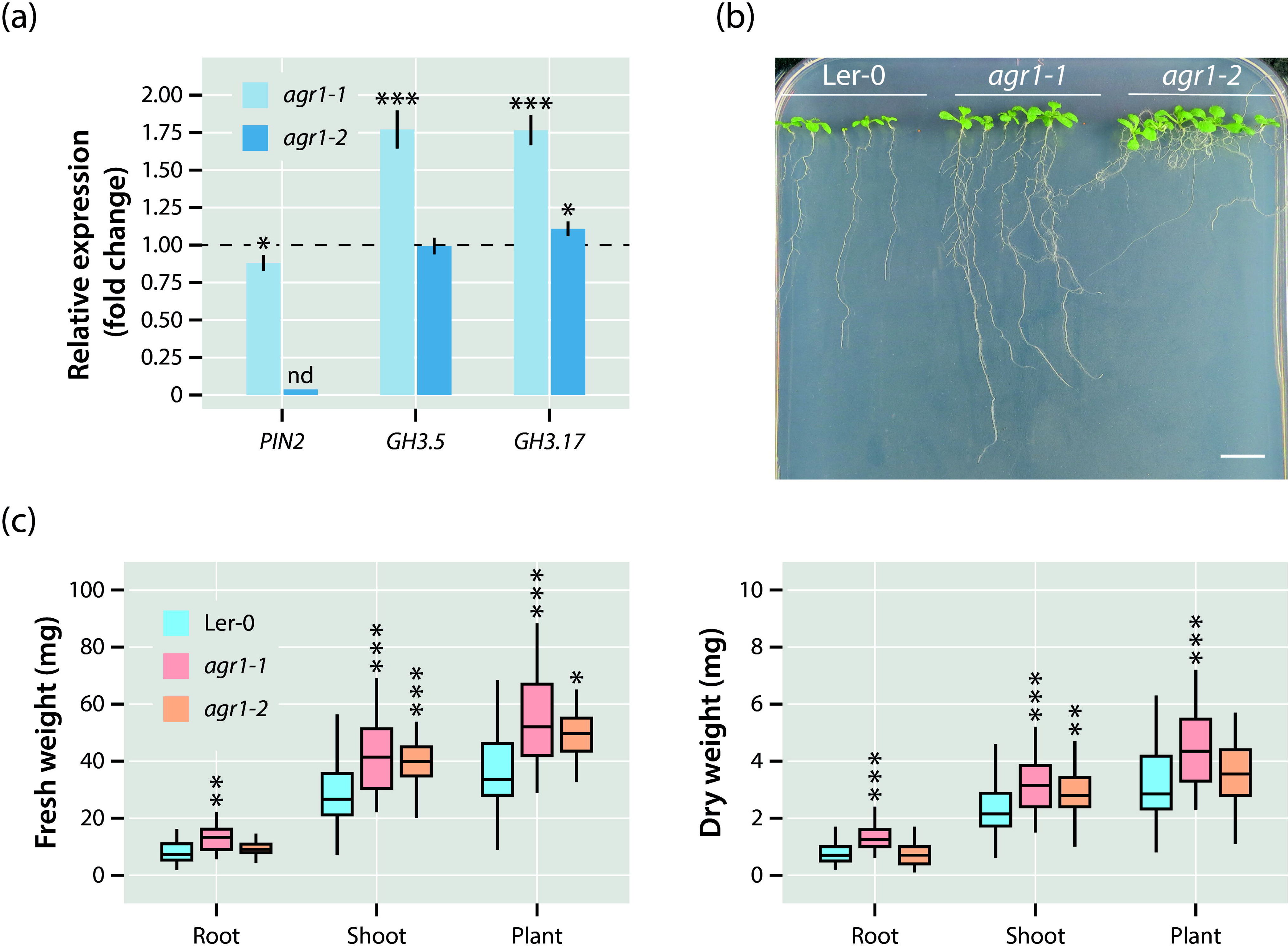
Transcriptional and physiological effects of a partial loss of *PIN2*. (**a**) Quantification of *PIN2*, *GH3.5*, and *GH3.17* expression in the mutants *agr1-1* and *agr1-2* relative to the Ler-0 wild type (dashed line) by qRT-PCR. The bars show means of n = 3 independent measurements. Asterisks mark the experiments with significantly altered gene expressions. Student’s *t*-test: *p ≤ 0.05, ***p ≤ 0.001. (**b**) Phenotype of the used Arabidopsis genotypes, Ler-0, *agr1-1*, and *agr1-2* grown for 10 days on vertical 0.5× MS plates. Scale bar = 1 cm. (**c**) Fresh and dry weight measurements of the tested Arabidopsis genotypes. The box plots show the median, quartiles, and extremes of the compared data sets (n = 32). Asterisks indicate significant differences between the Ler-0 control and the corresponding *agr1-1* and *agr1-2* samples. Student’s *t*-test: *p ≤ 0.05, **p ≤ 0.01, ***p ≤ 0.001.

As depicted in **Figure 9a**, in addition to the functional mutation, *PIN2* expression is approximately 10-15% lower in *agr1-1* compared to wild type. On the contrary, the expression of *PIN2* was not detectable in *agr1-2*. While the functional loss of PIN2 in *agr1-2* had hardly any impact on *GH3.5* and *GH3.17* expression, the two genes were found to be considerably induced in *agr1-1*, suggesting that subtle changes in PIN2 abundance and/or functionality can trigger *GH3.5* and *GH3.17* induction, while an entire loss of PIN2 does not translate into an induction of the genes. Interestingly, as shown in **Figure 9b,c**, the partial loss of PIN2 in *agr1- 1* also led to a consistent increased of root and shoot biomass, both in terms of fresh and dry weight. The effect of the *agr1-2* mutation had less impact on the plant biomass and was largely restricted to an increase in shoot biomass. Taken together, the alteration of local auxin contents in the root tips due to the repression of *PIN2* is likely sufficient to boost plant growth and to induce the expression of specific *GH3* genes that keep the local auxin accumulation under control.

## 4. Discussion

The growth promoting effect of the root-colonizing endophyte *S. indica* on a wide variety of host plants, including canola, sweet potato, barley, rice and the dicot model plant *A. thaliana,* is widely documented in the literature (Bagheri et al., 2013; Li et al., 2021; Pérez-Alonso et al., 2022; Su et al., 2017; Waller et al., 2005). Especially the substantial stimulation of root growth of *S. indica*-infected plants raised a lot of attention because root system architecture is intimately linked to plant productivity and stress tolerance (Comas et al., 2013; Khan et al., 2016; Lynch, 1995). Root growth and development are closely coordinated through the sophisticated crosstalk of plant hormones. In this context, auxin, ethylene, cytokinin, abscisic acid, gibberellin, and brassinosteroids collaborate in a complex regulatory network involving synergistic or antagonistic interactions of the contributing signaling molecules. Within this network, the directional transport of auxin through the root provides crucial positional information that is important for proper root development (Vanneste & Friml, 2009). The distribution of auxin along the developing root is vital for its proper growth and development, including the formation of lateral roots. In Arabidopsis roots, patterning is achieved through the formation of a local auxin maximum at the root tip (Sabatini et al., 1999). The establishment and coordination of auxin maxima in tips of primary and lateral roots is the product of the complex interplay between auxin transport (Petrášek & Friml, 2009) and local auxin biosynthesis (Ikeda et al., 2009; Zhao, 2010).

Several studies, employing transcriptomics and metabolomics approaches, have already addressed the question of whether *S. indica* alters auxin biosynthesis in its host plants to trigger the considerable induction of root growth observed (Hilbert et al., 2012; Lahrmann et al., 2015). However, none of these studies reported consistent induction of auxin biosynthesis-related genes in plants infected with *S. indica,* but only provided evidence for the transcriptional activation of pathways that promote the formation of L-Trp derivatives, including indole glucosinolates, camalexin, and indole-3-carboxylic acid, which play important roles in limiting endophyte growth within the host plant (Lahrmann et al., 2015; Nongbri et al., 2012). However, both auxin content and auxin signaling have been shown to increase in the early infection phase, leading to the assumption that *S. indica* provides IAA during the initial infection phase through its own biosynthetic pathway (Hilbert et al., 2012; Meents et al., 2019; Sirrenberg et al., 2007; Vadassery et al., 2008). In fact, the observations made in this study confirmed the previously published data (**Figure 1**). However, our study complementarily detected a significant increase in IAA-Asp levels at later stages of the interaction (**Figure 2a**). Many auxin conjugates, including IAA-Ala, indole-3-acetyl-L-leucine and indole-3-acetyl-L-phenylalanine, serve as temporal storage forms for auxin, as they can be hydrolyzed to release physiologically active free IAA by specific IAA amino acid conjugate hydrolases in Arabidopsis (LeClere et al., 2002; Rampey et al., 2004). The conjugation of IAA with L-Asp and L-Glu, however, appears to be irreversible and, thus, marks a first step in auxin degradation (Östin et al., 1998; Rampey et al., 2004). Our observation of the involvement of auxin conjugating IAA-amido synthetases of the GRETCHEN HAGEN 3 (GH3) family and increased auxin signaling activities in the early stages of infection are in agreement with the known formation of IAA by *S. indica*. At 3 dpi, auxin contents were detected to be substantially higher in plants infected with the fungus. However, over time, the free auxin content decreased, no longer showing considerable differences to the controls, while auxin conjugation apparently increased. An increase in auxin conjugation in parallel with a decrease in free auxin contents is not necessarily in accordance with the high auxin signaling activity detected in plants infected with the fungus at 6 dpi (**Figure 2c,d**). However, these results suggested the formation of local auxin maxima that can derive from altered auxin transport activities and sparked our interest in further investigating the role of GH3 enzymes in the symbiosis between Arabidopsis and *S. indica*. Interestingly, auxin signaling showed no considerable difference between mock- and fungus-infected samples at 3 dpi. This could possibly be due to the transient alkalinization of the rhizosphere in *S. indica*-infected roots, as this would hamper the diffusion of IAA derived from the fungus into the root cells (Lanza et al., 2019).

Until now, there has been only very limited information on the role of GH3 enzymes in plant–fungus interactions. A previous work reported the induction of an Arabidopsis *GH3.6* homolog gene (Potri.011G129700) in *Populus trichocarpa* inoculated with either *Mortierella elongata* or *Ilyonectria europaea* (Liao et al., 2019), while another study highlighted the induction of an Arabidopsis *GH3.17* homolog gene by the endophytic fungus *Chaetomium cupreum* in roots of *Eucalyptus globulus* (Ortiz et al., 2019). Furthermore, a study on the role of cytokinin and auxin in the interaction between Arabidopsis and *S. indica* demonstrated that the high auxin phenotype of the *sur1-1* mutant can be rescued by colonization with the fungus (Vadassery et al., 2008). Nonetheless, the latter study only considered changes in total conjugated auxin contents, which did not show significant alterations in *S. indica*-infected plants. This left room to speculate that auxin degradation or a missing limitation of root colonization could be involved in altering the *sur1-1* mutant phenotype.

To further address the question of the involvement of auxin conjugation in Arabidopsis–*S. indica* interactions, we used an alternative high auxin mutant, YUC9ox, which is known to have an approximately 2.5 times higher IAA content (Hentrich et al., 2013). Infection of YUC9ox with *S. indica* (**Figure 3a**) clearly confirmed a considerable impact of the fungus on the strong high auxin phenotype of the mutant. RNA-seq analysis of mock and *S. indica* infected wt and YUC9ox plants provided evidence for the induction of a small number of genes that respond to high auxin (YUC9ox versus Col-0 up, –E) and to the presence of the endophyte (YUC9ox versus Col-0 up, +E) (**Figure 3d**). The subsequent GO analysis of the 402 selected genes highlighted the overrepresentation of 33 auxin response-related genes, including several *GH3* genes (**Figure 3e**). This finding largely supported our hypothesis of a crucial role for GH3 enzymes in the plant–fungus consortium investigated. Directed analysis of *GH3* genes in *S. indica*-versus mock-infected Col-0 seedlings supported our hypothesis and focused our interest on the three *GH3* genes *GH3.4*, *GH3.5*, and *GH3.17*. The genes responded rapidly to fungus infection of the roots (**Figure 4**). To further confirm these indications, we took a reverse genetics approach and tested the growth promoting effect of *S. indica* on roots of Arabidopsis wt and *gh3* mutants. Our study clearly highlighted the involvement of GH3.5 and GH3.17 (**Figure 5a, Supporting Information: Image 2**). While the *gh3.4* mutant showed no considerable differences to the wild-type control, the two other individual mutants are clearly compromised in the interaction with the fungus, as the growth promoting effect was largely absent.

GH3.5 and GH3.17 were recently shown to contribute to the control of root elongation (Guo et al., 2022). This added to a previous report on GH3.17, demonstrating its involvement in hypocotyl elongation during the shade avoidance reaction (Zheng et al., 2016). A triple regulatory role has been attributed to GH3.5, which conjugates IAA and, in addition, salicylic acid (SA) and jasmonic acid (JA) to modulate auxin and pathogen responses (Gutierrez et al., 2012; Westfall et al., 2016; Zhang et al., 2007). This triple function featured GH3.5 as an important mediator in the allocation of metabolic resources to establish improved resistance traits to pathogens (Park et al., 2007). Intriguingly, the knockout of all eight members of group II of *GH3* genes only resulted in an increased density of the lateral roots, without affecting the overall length of the primary roots. Furthermore, the octuple mutant was shown to be more salt- and drought-tolerant, presumably through the increased IAA content in the mutant (Casanova-Sáez et al., 2022).

To explore the role of GH3.5 and GH3.17 in more detail, we addressed the question if the induction of the two genes observed in transcriptomics experiments (**Figure 4**) is locally restricted or detectable in all root areas penetrated by the fungus. This was particularly interesting, because previous studies demonstrated that the colonization of Arabidopsis roots with *S. indica* occurs in the elongation and maturation zones of the root (Jacobs et al., 2011), while *GH3.5* and *GH3.17* were shown to be predominantly expressed at the tips of the root (Pierdonati et al., 2019). Confocal laser scanning microscopy allowed us to monitor the induction profiles of *GH3.5* and *GH3.17* at the cellular level by employing corresponding promoter–reporter constructs. The obtained results explicitly pointed to an induction of *GH3.5* and *GH3.17* in the tips of primary and lateral roots (**Figure 7, Supporting Information:Image 4**). The critical role of auxin conjugation in the establishment of the beneficial plant–fungus interaction was further strengthened by the analysis of auxin conjugate hydrolase overexpressing *35S::IAR3* mutant lines (**Figure 6**). The apparent lack of growth promotion in line 1.3 and the considerably reduced growth promotion in the weaker 6.2 line compared to the wild-type control clearly highlights the importance of a tightly controlled cellular auxin homeostasis through the local induction of *GH3.5* and *GH3.17* in the root tips over the course of the infection of Arabidopsis with *S. indica*.

As already mentioned, root tips show no substantial fungus colonization. Therefore, the question of how the expression of *GH3.5* and *GH3.17* can be induced by the fungus remained to be answered. Taking into account the persistently high auxin signaling activities in roots infected with the fungus, we concluded that the induction of the two *GH3* genes is likely a secondary effect that involves increased contents of auxin at the root tips through an adjustment of auxin transport, which, in turn, is known to quickly stimulate the production of GH3 enzymes (Hagen & Guilfoyle, 2002). In fact, our experiments revealed the transcriptional repression of *PIN2* in fungus infected root areas (**Figure 8**), which can eventually result in the local accumulation of auxin in the root tips. Interestingly, alteration of auxin transport appears to be a more general theme in plant–microbe interactions. A recent work already reported the role of the transcriptional modification of auxin exporters in the symbiosis between *Bradyrhizobium japonicum* and *A. thaliana* (Schroeder et al., 2022). However, in contrast to the significant repression of *PIN2* in *S. indica* infections, the plant growth promoting effect of the bacterium appears to be mainly associated with the induction of the expression of *PIN3*, *PIN7*, and *ABCB19*.

The notion that the expression of *GH3.5* and *GH3.17* is the consequence of changes in auxin transport in the root was confirmed by analyzing gene expression levels in two independent *PIN2* mutants, *agr1-1* and *agr1-2* (Bell & Maher, 1990; Chen et al., 1998). However, it should be noted that only the modest reduction of *PIN2* expression and the less severe point mutation in *agr1-1* significantly triggered the expression of *GH3.5* and *GH3.17*, thereby mimicking the situation roots co-cultivated with *S. indica*, while a complete loss of *PIN2*, as in *agr1-2*, had no considerable effect on the expression level of the two genes (**Figure 9**). The genetically forced auxin accumulation in maize root tips through the overexpression of *PIN1* already underlined a significant impact of altered auxin maxima at the tip of the roots on plant growth (Li et al., 2018). This led us to conclude that possibly the subtle repression of *PIN2* over the course of an infection with *S. indica* is sufficient to fine-tune auxin contents in root tips, which eventually may provoke the induction of root growth and, subsequently, also lead to an increased biomass production in the shoot, most likely due to an improved assimilation of nutrients. When the biomass production of the *agr1-1* and *agr1-2* mutants was compared to the values obtained for similarly grown wild-type Ler-0 plants, a significant increase in plant biomass in *agr1-1*, but not in *agr1-2* was detected. In the latter mutant, only the shoot fresh and dry weight appeared to be higher than in the control plants.

In summary, our work provides for the first time evidence for an essential role of auxin transport alterations that appear to be involved in triggering the growth promoting effect of *S. indica* in its host plant. Infection of Arabidopsis with the root colonizing fungus activates auxin signaling and represses the expression of the auxin exporter gene *PIN2*, resulting in reduced basipetal auxin redirection and, thus, accumulation of auxin at the tips of the roots. However, our experiments also pointed out that the adjustment of auxin transport in the symbiotic system studied was very delicate. A complete loss of PIN2, as in the *agr1-2* mutant, has only a very reduced impact on plant biomass production. In contrast, our results clearly demonstrate that a subtle reduction in the abundance or functionality of PIN2, as in case of *S. indica*-infected roots and in the *agr1-1* mutant, respectively, significantly promotes plant biomass production. Furthermore, our work pinpoints the crucial role of the interplay between the formation of local auxin maxima at the root tips and its local conjugation through the induction of specific GH3 IAA amido-synthetase genes, to keep auxin accumulation under control. Therefore, our work provides novel information on the molecular mechanism that contributes to the adjustment of the local auxin distribution in roots that is involved in the *S. indica*-stimulated alteration of the root morphology and contributes to a greater understanding of the symbiotic plant–fungus interaction.

## 6. Acknowledgements

The authors thank Paul E. Staswick (University of Nebraska-Lincoln, USA) for kindly providing the *gh3* mutant lines and to Miguel Ángel Moreno-Risueño and Mary Paz González-García (Centro de Biotecnología y Genómica de Plantas, Spain) as well as to Riccardo Di Mambro (University of Pisa, Italy) for sharing *DR5::Luc*, *pPIN2::PIN2-GFP*, and *pGH3.5::3*×*YFP* as well as *pGH3.17::3*×*YFP* seeds, respectively, with us. Furthermore, the authors are grateful to Mar González Ceballos (CBGP) and Freia Benade (TU Dresden) for their excellent technical support.

## 8. Authors contributions

SP, RO, JLM, JVC, AK and HS conceived and planned the study. AGOV, EK, MKP, MMPA, SSS, TK, MK, YT, PR, LMQ, SB, AS, BSS, NB, and SP performed the experiments and collected the data. AGOV, JVC, and SP analyzed and interpreted the data; SP wrote the paper with significant input from all other authors.

## 9. Data Availability Statement

All data supporting the conclusions of this study are present in the paper and/or the Supporting Information. The GEO accession numbers for the raw and processed RNA-seq data used in this study are GSE240683 and GSE241902.

## 10. Short legends for Supporting Information

Supporting Information:Data Sheet 1: Primers used for genotyping, cloning and expression analysis.

Supporting Information:Image 1: GO chord chart of DEGs (*S. indica* vs mock) at 10 dpi.

Supporting Information:Image 2: Results of UHPLC-MS/MS analysis of *S. indica* and mock treated Arabidopsis wild-type seedlings.

Supporting Information:Image 3: Representative images of the growth phenotype of *gh3* and *35S::IAR3* mutants.

Supporting Information:Image 4: *S. indica*-triggered induction of *GH3.5* and *GH3.17* expression in *pGH3.5::3*×*YFP* and *pGH3.17::3*×*YFP* (fluorescence channel).

Supporting Information:Image 5: *GH3.5* and *GH3.17* expression in mock- and *S. indica*-infected *pGH3.5::Luc* and *pGH3.17::Luc* lines.

## References

Bagheri, A., Saadatmand, S., Niknam, V., Nejadsatari, T., & Babaeizad, V. (2013). Effect of Endophytic Fungus, *Piriformospora Indica*, on Growth and Activity of Antioxidant Enzymes of Rice (*Oryza Sativa* L.) Under Salinity Stress. International Journal of Advanced Biological and Biomedical Research, 1, 1337–1350. https://www.ijabbr.com/article_7908.html

Barlier, I., Kowalczyk, M., Marchant, A., Ljung, K., Bhalerao, R., Bennett, M., Sandberg, G., & Bellini, C. (2000). The *SUR2* gene of *Arabidopsis thaliana* encodes the cytochrome P450 CYP83B1, a modulator of auxin homeostasis. Proceedings of the National Academy of Sciences USA, 97(26), 14819–14824. 10.1073/pnas.260502697

Bastías, D. A., Balestrini, R., Pollmann, S., & Gundel, P. E. (2022). Environmental interference of plant-microbe interactions. Plant, Cell & Environment, 45(12), 3387–3398. 10.1111/pce.14455

Bell, C. J., & Maher, E. P. (1990). Mutants of *Arabidopsis thaliana* with abnormal gravitropic responses. Molecular and General Genetics MGG, 220(2), 289–293. 10.1007/BF00260496

Böttcher, C., Westphal, L., Schmotz, C., Prade, E., Scheel, D., & Glawischnig, E. (2009). The multifunctional enzyme CYP71B15 (PHYTOALEXIN DEFICIENT3) converts cysteine-indole-3-acetonitrile to camalexin in the indole-3-acetonitrile metabolic network of *Arabidopsis thaliana*. Plant Cell, 21(6), 1830–1845. 10.1105/tpc.109.066670

Brumos, J., Robles, L. M., Yun, J., Vu, T. C., Jackson, S., Alonso, J. M., & Stepanova, A. N. (2018). Local Auxin Biosynthesis Is a Key Regulator of Plant Development. Developmental Cell, 47(3), 306–318 e305. 10.1016/j.devcel.2018.09.022

Casanova-Sáez, R., Mateo-Bonmatí, E., Šimura, J., Pěnčík, A., Novák, O., Staswick, P., & Ljung, K. (2022). Inactivation of the entire Arabidopsis group II GH3s confers tolerance to salinity and water deficit. New Phytologist, 235(1), 263–275. 10.1111/nph.18114

Charura, N. M., Llamas, E., De Quattro, C., Vilchez, D., Nowack, M. K., & Zuccaro, A. (2023). Root cap cell corpse clearance limits microbial colonization in *Arabidopsis thaliana*. bioRxiv, 2023.2002.2003.526420. 10.1101/2023.02.03.526420

Chen, R., Hilson, P., Sedbrook, J., Rosen, E., Caspar, T., & Masson, P. H. (1998). The *Arabidopsis thaliana AGRAVITROPIC 1* gene encodes a component of the polar-auxin-transport efflux carrier. Proceedings of the National Academy of Sciences USA, 95(25), 15112–15117. 10.1073/pnas.95.25.15112

Clough, S. J., & Bent, A. F. (1998). Floral dip: a simplified method for *Agrobacterium*-mediated transformation of *Arabidopsis thaliana*. Plant Journal, 16(6), 735–743. 10.1046/j.1365-313x.1998.00343.x

Comas, L., Becker, S., Cruz, V. M., Byrne, P. F., & Dierig, D. A. (2013). Root traits contributing to plant productivity under drought. Frontiers in Plant Science, 4, 442. 10.3389/fpls.2013.00442

Curtis, M. D., & Grossniklaus, U. (2003). A Gateway Cloning Vector Set for High-Throughput Functional Analysis of Genes in Planta. Plant Physiology, 133(2), 462–469. 10.1104/pp.103.027979

Czechowski, T., Stitt, M., Altmann, T., Udvardi, M. K., & Scheible, W. R. (2005). Genome-wide identification and testing of superior reference genes for transcript normalization in Arabidopsis. Plant Physiology, 139(1), 5–17. 10.1104/pp.105.063743

Davies, R. T., Goetz, D. H., Lasswell, J., Anderson, M. N., & Bartel, B. (1999). *IAR3* encodes an auxin conjugate hydrolase from Arabidopsis. Plant Cell, 11(3), 365–376. 10.1105/tpc.11.3.365

Fröschel, C., Komorek, J., Attard, A., Marsell, A., López-Arboleda, W. A., Le Berre, J., Wolf, E., Geldner, N., Waller, F., Korte, A., & Dröge-Laser, W. (2021). Plant roots employ cell-layer-specific programs to respond to pathogenic and beneficial microbes. Cell Host & Microbe, 29(2), 299–310.e297. 10.1016/j.chom.2020.11.014

Galkovskyi, T., Mileyko, Y., Bucksch, A., Moore, B., Symonova, O., Price, C. A., Topp, C. N., Iyer-Pascuzzi, A. S., Zurek, P. R., Fang, S., Harer, J., Benfey, P. N., & Weitz, J. S. (2012). GiA Roots: software for the high throughput analysis of plant root system architecture. BMC Plant Biology, 12(1), 116. 10.1186/1471-2229-12-116

Guo, R., Hu, Y., Aoi, Y., Hira, H., Ge, C., Dai, X., Kasahara, H., & Zhao, Y. (2022). Local conjugation of auxin by the GH3 amido synthetases is required for normal development of roots and flowers in Arabidopsis. Biochemical and Biophysical Research Communications, 589, 16–22. 10.1016/j.bbrc.2021.11.109

Gutierrez, L., Mongelard, G., Flokova, K., Pacurar, D. I., Novak, O., Staswick, P., Kowalczyk, M., Pacurar, M., Demailly, H., Geiss, G., & Bellini, C. (2012). Auxin controls Arabidopsis adventitious root initiation by regulating jasmonic acid homeostasis. Plant Cell, 24(6), 2515–2527. 10.1105/tpc.112.099119

Hagen, G., & Guilfoyle, T. (2002). Auxin-responsive gene expression: genes, promoters and regulatory factors. Plant Molecular Biology, 49(3), 373–385. 10.1023/A:1015207114117

Hentrich, M., Böttcher, C., Düchting, P., Cheng, Y., Zhao, Y., Berkowitz, O., Masle, J., Medina, J., & Pollmann, S. (2013). The jasmonic acid signaling pathway is linked to auxin homeostasis through the modulation of *YUCCA8* and *YUCCA9* gene expression. Plant Journal, 74(4), 626–637. 10.1111/tpj.12152

Hilbert, M., Voll, L. M., Ding, Y., Hofmann, J., Sharma, M., & Zuccaro, A. (2012). Indole derivative production by the root endophyte *Piriformospora indica* is not required for growth promotion but for biotrophic colonization of barley roots. New Phytologist, 196(2), 520–534. 10.1111/j.1469-8137.2012.04275.x

Hosseini, F., Mosaddeghi, M. R., & Dexter, A. R. (2017). Effect of the fungus *Piriformospora indica* on physiological characteristics and root morphology of wheat under combined drought and mechanical stresses. Plant Physiology and Biochemistry, 118, 107–120. 10.1016/j.plaphy.2017.06.005

Ikeda, Y., Men, S., Fischer, U., Stepanova, A. N., Alonso, J. M., Ljung, K., & Grebe, M. (2009). Local auxin biosynthesis modulates gradient-directed planar polarity in Arabidopsis. Nature Cell Biology, 11(6), 731–738. 10.1038/ncb1879

Jacobs, S., Zechmann, B., Molitor, A., Trujillo, M., Petutschnig, E., Lipka, V., Kogel, K. H., & Schäfer, P. (2011). Broad-spectrum suppression of innate immunity is required for colonization of Arabidopsis roots by the fungus *Piriformospora indica*. Plant Physiology, 156(2), 726–740. 10.1104/pp.111.176446

Jahn, L., Mucha, S., Bergmann, S., Horn, C., Staswick, P., Steffens, B., Siemens, J., & Ludwig-Müller, J. (2013). The Clubroot Pathogen (*Plasmodiophora brassicae*) Influences Auxin Signaling to Regulate Auxin Homeostasis in Arabidopsis. Plants, 2(4), 726–749. https://www.mdpi.com/2223-7747/2/4/726

Jogawat, A., Vadassery, J., Verma, N., Oelmüller, R., Dua, M., Nevo, E., & Johri, A. K. (2016). PiHOG1, a stress regulator MAP kinase from the root endophyte fungus *Piriformospora indica*, confers salinity stress tolerance in rice plants. Scientific Reports, 6, 36765. 10.1038/srep36765

Johnson, J. M., Sherameti, I., Nongbri, P. L., & Oelmüller, R. (2013). Standardized Conditions to Study Beneficial and Nonbeneficial Traits in the *Piriformospora indica*/*Arabidopsis thaliana* Interaction. In A. Varma, G. Kost, & R. Oelmüller (Eds.), Piriformospora indica: Sebacinales and Their Biotechnological Applications (pp. 325–343). Springer Berlin Heidelberg. 10.1007/978-3-642-33802-1_20

Jost, R., Berkowitz, O., & Masle, J. (2007). Magnetic quantitative reverse transcription PCR: A high-throughput method for mRNA extraction and quantitative reverse transcription PCR. BioTechniques, 43(2), 206–211. 10.2144/000112534

Khan, M. A., Gemenet, D. C., & Villordon, A. (2016). Root System Architecture and Abiotic Stress Tolerance: Current Knowledge in Root and Tuber Crops. Frontiers in Plant Science, 7, 1584. 10.3389/fpls.2016.01584

Kojima, M., Kamada-Nobusada, T., Komatsu, H., Takei, K., Kuroha, T., Mizutani, M., Ashikari, M., Ueguchi-Tanaka, M., Matsuoka, M., Suzuki, K., & Sakakibara, H. (2009). Highly Sensitive and High-Throughput Analysis of Plant Hormones Using MS-Probe Modification and Liquid Chromatography–Tandem Mass Spectrometry: An Application for Hormone Profiling in Oryza sativa. Plant and Cell Physiology, 50(7), 1201–1214. 10.1093/pcp/pcp057

Lahrmann, U., Strehmel, N., Langen, G., Frerigmann, H., Leson, L., Ding, Y., Scheel, D., Herklotz, S., Hilbert, M., & Zuccaro, A. (2015). Mutualistic root endophytism is not associated with the reduction of saprotrophic traits and requires a noncompromised plant innate immunity. New Phytologist, 207(3), 841–857. 10.1111/nph.13411

Lanza, M., Haro, R., Conchillo, L. B., & Benito, B. (2019). The endophyte *Serendipita indica* reduces the sodium content of Arabidopsis plants exposed to salt stress: fungal ENA ATPases are expressed and regulated at high pH and during plant co-cultivation in salinity. Environmental Microbiology, 21(9), 3364–3378. 10.1111/1462-2920.14619

LeClere, S., Tellez, R., Rampey, R. A., Matsuda, S. P. T., & Bartel, B. (2002). Characterization of a Family of IAA-Amino Acid Conjugate Hydrolases from *Arabidopsis*. Journal of Biological Chemistry, 277(23), 20446–20452. 10.1074/jbc.M111955200

Li, Q., Kuo, Y.-W., Lin, K.-H., Huang, W., Deng, C., Yeh, K.-W., & Chen, S.-P. (2021). *Piriformospora indica* colonization increases the growth, development, and herbivory resistance of sweet potato (*Ipomoea batatas* L.). Plant Cell Reports, 40(2), 339–350. 10.1007/s00299-020-02636-7

Li, Z., Zhang, X., Zhao, Y., Li, Y., Zhang, G., Peng, Z., & Zhang, J. (2018). Enhancing auxin accumulation in maize root tips improves root growth and dwarfs plant height. Plant Biotechnology Journal, 16(1), 86–99. 10.1111/pbi.12751

Liao, H.-L., Bonito, G., Rojas, J. A., Hameed, K., Wu, S., Schadt, C. W., Labbé, J., Tuskan, G. A., Martin, F., Grigoriev, I. V., & Vilgalys, R. (2019). Fungal Endophytes of *Populus trichocarpa* Alter Host Phenotype, Gene Expression, and Rhizobiome Composition. Molecular Plant-Microbe Interactions, 32(7), 853–864. 10.1094/mpmi-05-18-0133-r

Livak, K. J., & Schmittgen, T. D. (2001). Analysis of relative gene expression data using real-time quantitative PCR and the 2^-ΔΔC(t)^ Method. Methods, 25(4), 402–408. 10.1006/meth.2001.1262

Lynch, J. (1995). Root Architecture and Plant Productivity. Plant Physiology, 109(1), 7–13. 10.1104/pp.109.1.7

Mateo-Bonmatí, E., Casanova-Sáez, R., Simura, J., & Ljung, K. (2021). Broadening the roles of UDP-glycosyltransferases in auxin homeostasis and plant development. New Phytologist, 232(2), 642–654. 10.1111/nph.17633

Meents, A. K., Furch, A. C. U., Almeida-Trapp, M., Özyürek, S., Scholz, S. S., Kirbis, A., Lenser, T., Theissen, G., Grabe, V., Hansson, B., Mithöfer, A., & Oelmüller, R. (2019). Beneficial and Pathogenic Arabidopsis Root-Interacting Fungi Differently Affect Auxin Levels and Responsive Genes During Early Infection. Frontiers in Microbiology, 10, 380. 10.3389/fmicb.2019.00380

Mellor, N., Band, L. R., Pěnčík, A., Novák, O., Rashed, A., Holman, T., Wilson, M. H., Voss, U., Bishopp, A., King, J. R., Ljung, K., Bennett, M. J., & Owen, M. R. (2016). Dynamic regulation of auxin oxidase and conjugating enzymes *AtDAO1* and *GH3* modulates auxin homeostasis. Proceedings of the National Academy of Sciences USA, 113(39), 11022–11027. 10.1073/pnas.1604458113

Mensah, R. A., Li, D., Liu, F., Tian, N., Sun, X., Hao, X., Lai, Z., & Cheng, C. (2020). Versatile *Piriformospora indica* and Its Potential Applications in Horticultural Crops. Horticultural Plant Journal, 6(2), 111–121. 10.1016/j.hpj.2020.01.002

Michniewicz, M., Brewer, P. B., & Friml, J. (2007). Polar auxin transport and asymmetric auxin distribution. The Arabidopsis book, 5, e0108–e0108. 10.1199/tab.0108

Moreno-Risueno, M. A., Van Norman, J. M., Moreno, A., Zhang, J., Ahnert, S. E., & Benfey, P. N. (2010). Oscillating Gene Expression Determines Competence for Periodic *Arabidopsis* Root Branching. Science, 329(5997), 1306–1311. 10.1126/science.1191937

Müller, A., Hillebrand, H., & Weiler, E. W. (1998). Indole-3-acetic acid is synthesized from L-tryptophan in roots of *Arabidopsis thaliana*. Planta, 206(3), 362–369. 10.1007/s004250050411

Müller, T. M., Böttcher, C., & Glawischnig, E. (2019). Dissection of the network of indolic defence compounds in *Arabidopsis thaliana* by multiple mutant analysis. Phytochemistry, 161, 11–20. 10.1016/j.phytochem.2019.01.009

Murashige, T., & Skoog, F. (1962). A revised medium for rapid growth and bio assays with tobacco tissue cultures. Physiologia Plantarum, 15(3), 473–497. 10.1111/j.1399-3054.1962.tb08052.x

Nakagawa, T., Suzuki, T., Murata, S., Nakamura, S., Hino, T., Maeo, K., Tabata, R., Kawai, T., Tanaka, K., Niwa, Y., Watanabe, Y., Nakamura, K., Kimura, T., & Ishiguro, S. (2007). Improved Gateway Binary Vectors: High-Performance Vectors for Creation of Fusion Constructs in Transgenic Analysis of Plants. Bioscience, Biotechnology, and Biochemistry, 71(8), 2095–2100. 10.1271/bbb.70216

Nongbri, P. L., Johnson, J. M., Sherameti, I., Glawischnig, E., Halkier, B. A., & Oelmuller, R. (2012). Indole-3-acetaldoxime-derived compounds restrict root colonization in the beneficial interaction between *Arabidopsis* roots and the endophyte *Piriformospora indica*. Molecular Plant-Microbe Interaction, 25(9), 1186–1197. 10.1094/MPMI-03-12-0071-R

Oñate-Sánchez, L., & Vicente-Carbajosa, J. (2008). DNA-free RNA isolation protocols for *Arabidopsis thaliana*, including seeds and siliques. BMC Research Notes, 1, 93. 10.1186/1756-0500-1-93

Ortiz, J., Soto, J., Fuentes, A., Herrera, H., Meneses, C., & Arriagada, C. (2019). The Endophytic Fungus *Chaetomium cupreum* Regulates Expression of Genes Involved in the Tolerance to Metals and Plant Growth Promotion in *Eucalyptus globulus* Roots. Microorganisms, 7(11), 490. 10.3390/microorganisms7110490

Östin, A., Kowalyczk, M., Bhalerao, R. P., & Sandberg, G. (1998). Metabolism of indole-3-acetic acid in Arabidopsis. Plant Physiology, 118(1), 285–296. 10.1104/pp.118.1.285

Park, J. E., Park, J. Y., Kim, Y. S., Staswick, P. E., Jeon, J., Yun, J., Kim, S. Y., Kim, J., Lee, Y. H., & Park, C. M. (2007). GH3-mediated auxin homeostasis links growth regulation with stress adaptation response in *Arabidopsis*. Journal of Biological Chemistry, 282(13), 10036–10046. 10.1074/jbc.M610524200

Pérez-Alonso, M. M., Guerrero-Galán, C., González Ortega-Villaizan, A., Ortiz-García, P., Scholz, S. S., Ramos, P., Sakakibara, H., Kiba, T., Ludwig-Müller, J., Krapp, A., Oelmüller, R., Vicente-Carbajosa, J., & Pollmann, S. (2022). The calcium sensor CBL7 is required for *Serendipita indica*-induced growth stimulation in *Arabidopsis thaliana*, controlling defense against the endophyte and K^+^ homoeostasis in the symbiosis. *Plant*, Cell & Environment, 45(11), 3367–3382. 10.1111/pce.14420

Pérez-Alonso, M. M., Guerrero-Galán, C., Scholz, S. S., Kiba, T., Sakakibara, H., Ludwig-Müller, J., Krapp, A., Oelmüller, R., Vicente-Carbajosa, J., & Pollmann, S. (2020). Harnessing symbiotic plant-fungus interactions to unleash hidden forces from extreme plant ecosystems. Journal of Experimental Botany, 71(13), 3865–3877. 10.1093/jxb/eraa040

Pérez-Alonso, M. M., Ortiz-García, P., Moya-Cuevas, J., Lehmann, T., Sánchez-Parra, B., Björk, R. G., Karim, S., Amirjani, M. R., Aronsson, H., Wilkinson, M. D., & Pollmann, S. (2021b). Endogenous indole-3-acetamide levels contribute to the crosstalk between auxin and abscisic acid, and trigger plant stress responses in *Arabidopsis thaliana*. Journal of Experimental Botany, 72(2), 459–475. 10.1093/jxb/eraa485

Pérez-Alonso, M. M., Ortiz-García, P., Moya-Cuevas, J., & Pollmann, S. (2021a). Mass Spectrometric Monitoring of Plant Hormone Cross Talk During Biotic Stress Responses in Potato (*Solanum tuberosum* L.). Methods in Molecular Biology, 2354, 143–154. 10.1007/978-1-0716-1609-3_7

Peškan-Berghöfer, T., Shahollari, B., Giong, P. H., Hehl, S., Markert, C., Blanke, V., Kost, G., Varma, A., & Oelmüller, R. (2004). Association of *Piriformospora indica* with *Arabidopsis thaliana* roots represents a novel system to study beneficial plant– microbe interactions and involves early plant protein modifications in the endoplasmic reticulum and at the plasma membrane. Physiologia Plantarum, 122(4), 465–477. 10.1111/j.1399-3054.2004.00424.x

Petrášek, J., & Friml, J. (2009). Auxin transport routes in plant development. Development, 136(16), 2675–2688. 10.1242/dev.030353

Pierdonati, E., Unterholzner, S. J., Salvi, E., Svolacchia, N., Bertolotti, G., Dello Ioio, R., Sabatini, S., & Di Mambro, R. (2019). Cytokinin-Dependent Control of GH3 Group II Family Genes in the Arabidopsis Root. Plants, 8(4). 10.3390/plants8040094

Porco, S., Pěnčík, A., Rashed, A., Voß, U., Casanova-Sáez, R., Bishopp, A., Golebiowska, A., Bhosale, R., Swarup, R., Swarup, K., Peňáková, P., Novák, O., Staswick, P., Hedden, P., Phillips, A. L., Vissenberg, K., Bennett, M. J., & Ljung, K. (2016). Dioxygenase-encoding *AtDAO1* gene controls IAA oxidation and homeostasis in *Arabidopsis*. Proceedings of the National Academy of Sciences USA, 113(39), 11016–11021. 10.1073/pnas.1604375113

Rampey, R. A., LeClere, S., Kowalczyk, M., Ljung, K., Sandberg, G., & Bartel, B. (2004). A family of auxin-conjugate hydrolases that contributes to free indole-3-acetic acid levels during Arabidopsis germination. Plant Physiology, 135(2), 978–988. 10.1104/pp.104.039677

Raudvere, U., Kolberg, L., Kuzmin, I., Arak, T., Adler, P., Peterson, H., & Vilo, J. (2019). g:Profiler: a web server for functional enrichment analysis and conversions of gene lists. Nucleic Acids Research, 47(W1), W191–W198. 10.1093/nar/gkz369

Rodríguez-Navarro, A., & Ramos, J. (1984). Dual system for potassium transport in *Saccharomyces cerevisiae*. Journal of Bacteriology, 159(3), 940–945. 10.1128/jb.159.3.940-945.1984

Roychoudhry, S., & Kepinski, S. (2022). Auxin in Root Development. Cold Spring Harbor Perspectives in Biology, 14(4), a039933. 10.1101/cshperspect.a039933

Sabatini, S., Beis, D., Wolkenfelt, H., Murfett, J., Guilfoyle, T., Malamy, J., Benfey, P. N., Leyser, O., Bechtold, N., Weisbeek, P., & Scheres, B. (1999). An Auxin-Dependent Distal Organizer of Pattern and Polarity in the *Arabidopsis* Root. Cell, 99(5), 463–472. 10.1016/S0092-8674(00)81535-4

Schroeder, M. M., Gomez, M. Y., McLain, N., & Gachomo, E. W. (2022). *Bradyrhizobium japonicum* IRAT FA3 Alters *Arabidopsis thaliana* Root Architecture via Regulation of Auxin Efflux Transporters *PIN2*, *PIN3*, *PIN7*, and *ABCB19*. Molecular Plant-Microbe Interaction, 35(3), 215–229. 10.1094/MPMI-05-21-0118-R

Shinozaki, Y., Hao, S., Kojima, M., Sakakibara, H., Ozeki-Iida, Y., Zheng, Y., Fei, Z., Zhong, S., Giovannoni, J. J., Rose, J. K. C., Okabe, Y., Heta, Y., Ezura, H., & Ariizumi, T. (2015). Ethylene suppresses tomato (*Solanum lycopersicum*) fruit set through modification of gibberellin metabolism. Plant Journal, 83(2), 237–251. 10.1111/tpj.12882

Sirrenberg, A., Gobel, C., Grond, S., Czempinski, N., Ratzinger, A., Karlovsky, P., Santos, P., Feussner, I., & Pawlowski, K. (2007). *Piriformospora indica* affects plant growth by auxin production. Physiologia Plantarum, 131(4), 581–589. 10.1111/j.1399-3054.2007.00983.x

Staswick, P. E., Serban, B., Rowe, M., Tiryaki, I., Maldonado, M. T., Maldonado, M. C., & Suza, W. (2005). Characterization of an Arabidopsis enzyme family that conjugates amino acids to indole-3-acetic acid. Plant Cell, 17(2), 616–627. 10.1105/tpc.104.026690

Staswick, P. E., Tiryaki, I., & Rowe, M. L. (2002). Jasmonate response locus *JAR1* and several related Arabidopsis genes encode enzymes of the firefly luciferase superfamily that show activity on jasmonic, salicylic, and indole-3-acetic acids in an assay for adenylation. Plant Cell, 14(6), 1405–14015. 10.1105/tpc.000885

Stepanova, A. N., Robertson-Hoyt, J., Yun, J., Benavente, L. M., Xie, D. Y., Dolezal, K., Schlereth, A., Jürgens, G., & Alonso, J. M. (2008). TAA1-mediated auxin biosynthesis is essential for hormone crosstalk and plant development. Cell, 133(1), 177–191. 10.1016/j.cell.2008.01.047

Su, Z., Wang, T., Shrivastava, N., Chen, Y., Liu, X., Sun, C., Yin, Y., Gao, Q., & Lou, B. (2017). *Piriformospora indica* promotes growth, seed yield and quality of *Brassica napus* L. Microbiological Research, 199, 29–39. 10.1016/j.micres.2017.02.006

Sun, C., Shao, Y., Vahabi, K., Lu, J., Bhattacharya, S., Dong, S., Yeh, K. W., Sherameti, I., Lou, B., Baldwin, I. T., & Oelmüller, R. (2014). The beneficial fungus *Piriformospora indica* protects Arabidopsis from *Verticillium dahliae* infection by downregulation plant defense responses. BMC Plant Biology, 14, 268. 10.1186/s12870-014-0268-5

Utsuno, K., Shikanai, T., Yamada, Y., & Hashimoto, T. (1998). *AGR*, an *Agravitropic* Locus of *Arabidopsis thaliana*, Encodes a Novel Membrane-Protein Family Member. Plant and Cell Physiology, 39(10), 1111–1118. 10.1093/oxfordjournals.pcp.a029310

Vadassery, J., Ritter, C., Venus, Y., Camehl, I., Varma, A., Shahollari, B., Novák, O., Strnad, M., Ludwig-Müller, J., & Oelmüller, R. (2008). The role of auxins and cytokinins in the mutualistic interaction between Arabidopsis and *Piriformospora indica*. Molecular Plant-Microbe Interaction, 21(10), 1371–1383. 10.1094/MPMI-21-10-1371

van Dijk, M., Morley, T., Rau, M. L., & Saghai, Y. (2021). A meta-analysis of projected global food demand and population at risk of hunger for the period 2010–2050. Nature Food, 2(7), 494–501. 10.1038/s43016-021-00322-9

Vanneste, S., & Friml, J. (2009). Auxin: a trigger for change in plant development. Cell, 136(6), 1005–1016. 10.1016/j.cell.2009.03.001

Varma, A., Savita, V., Sudha, Sahay, N., Butehorn, B., & Franken, P. (1999). *Piriformospora indica*, a cultivable plant-growth-promoting root endophyte. Applied and Environmental Microbiology, 65(6), 2741–2744. 10.1128/AEM.65.6.2741-2744.1999

Waller, F., Achatz, B., Baltruschat, H., Fodor, J., Becker, K., Fischer, M., Heier, T., Hückelhoven, R., Neumann, C., von Wettstein, D., Franken, P., & Kogel, K. H. (2005). The endophytic fungus *Piriformospora indica* reprograms barley to salt-stress tolerance, disease resistance, and higher yield. Proceedings of the National Academy of Sciences USA, 102(38), 13386–13391. 10.1073/pnas.0504423102

Weiss, M., Waller, F., Zuccaro, A., & Selosse, M. A. (2016). Sebacinales - one thousand and one interactions with land plants. New Phytologist, 211(1), 20–40. 10.1111/nph.13977

Westfall, C. S., Sherp, A. M., Zubieta, C., Alvarez, S., Schraft, E., Marcellin, R., Ramirez, L., & Jez, J. M. (2016). *Arabidopsis thaliana* GH3.5 acyl acid amido synthetase mediates metabolic crosstalk in auxin and salicylic acid homeostasis. Proceedings of the National Academy of Sciences USA, 113(48), 13917–13922. 10.1073/pnas.1612635113

Wojtaczka, P., Ciarkowska, A., Starzynska, E., & Ostrowski, M. (2022). The GH3 amidosynthetases family and their role in metabolic crosstalk modulation of plant signaling compounds. Phytochemistry, 194, 113039. 10.1016/j.phytochem.2021.113039

Xu, J., & Scheres, B. (2005). Dissection of Arabidopsis ADP-RIBOSYLATION FACTOR 1 Function in Epidermal Cell Polarity. Plant Cell, 17(2), 525–536. 10.1105/tpc.104.028449

Xu, L., Wu, C., Oelmüller, R., & Zhang, W. (2018). Role of Phytohormones in *Piriformospora indica*-Induced Growth Promotion and Stress Tolerance in Plants: More Questions Than Answers. Frontiers in Microbiology, 9, 1646. 10.3389/fmicb.2018.01646

Zhang, W., Wang, J., Xu, L., Wang, A., Huang, L., Du, H., Qiu, L., & Oelmüller, R. (2018). Drought stress responses in maize are diminished by *Piriformospora indica*. Plant Signaling & Behavior, 13(1), e1414121. 10.1080/15592324.2017.1414121

Zhang, Z., Li, Q., Li, Z., Staswick, P. E., Wang, M., Zhu, Y., & He, Z. (2007). Dual regulation role of *GH3.5* in salicylic acid and auxin signaling during Arabidopsis-*Pseudomonas syringae* interaction. Plant Physiology, 145(2), 450–464. 10.1104/pp.107.106021

Zhang, Z., Wang, M., Li, Z., Li, Q., & He, Z. (2008). Arabidopsis GH3.5 regulates salicylic acid-dependent and both NPR1-dependent and independent defense responses. Plant Signaling & Behavior, 3(8), 537–542. 10.4161/psb.3.8.5748

Zhao, T., & Wang, Z. (2022). GraphBio: A shiny web app to easily perform popular visualization analysis for omics data [Original Research]. Frontiers in Genetics, 13. 10.3389/fgene.2022.957317

Zhao, Y. (2010). Auxin biosynthesis and its role in plant development. Annual Review of Plant Biology, 61, 49–64. 10.1146/annurev-arplant-042809-112308

Zheng, Z., Guo, Y., Novák, O., Chen, W., Ljung, K., Noel, J. P., & Chory, J. (2016). Local auxin metabolism regulates environment-induced hypocotyl elongation. Nature Plants, 2(4), 16025. 10.1038/nplants.2016.25

Zhou, Y., Zhou, B., Pache, L., Chang, M., Khodabakhshi, A. H., Tanaseichuk, O., Benner, C., & Chanda, S. K. (2019). Metascape provides a biologist-oriented resource for the analysis of systems-level datasets. Nature Communications, 10(1), 1523. 10.1038/s41467-019-09234-6

Zuccaro, A., Lahrmann, U., Guldener, U., Langen, G., Pfiffi, S., Biedenkopf, D., Wong, P., Samans, B., Grimm, C., Basiewicz, M., Murat, C., Martin, F., & Kogel, K. H. (2011). Endophytic life strategies decoded by genome and transcriptome analyses of the mutualistic root symbiont *Piriformospora indica*. PLoS Pathogens, 7(10), e1002290. 10.1371/journal.ppat.1002290

